# Polyamine homeostasis *in vivo* relies primarily on transport

**DOI:** 10.1101/2025.10.02.680054

**Authors:** Cindy Chang, Ankur Jain

## Abstract

Polyamines are essential metabolites present in all cells, but their regulation *in vivo* remains poorly understood. Little is known about whether tissues maintain distinct polyamine setpoints, how these setpoints are established, or whether such differences underlie selective vulnerability in disease. Here, we applied single-cell polyamine measurements in living *Caenorhabditis elegans* to map tissue polyamine levels and their regulatory dependencies. Three principles emerge: (i) Across differentiated tissues, steady-state polyamine pools are maintained primarily by transporter-mediated import rather than *de novo* synthesis. (ii) The intestine functions as a systemic regulator: perturbing intestinal polyamines affects organism-wide levels, and intestine-specific rescue restores systemic balance. (iii) Neurons maintain markedly low polyamine pools and undergo subtype-specific developmental reprogramming, switching polyamine acquisition strategies as they mature. These findings define core principles of tissue-specific polyamine regulation *in vivo* and provide a framework for developing therapeutic strategies to restore polyamine balance.

## Introduction

Polyamines are a class of small aliphatic metabolites containing two or more amine groups. These molecules are present in all domains of life and are maintained at near-millimolar concentrations in cells ^1–3^. The amino groups in polyamines are protonated at physiological pH, conferring a strong positive charge that enables electrostatic interactions with negatively-charged macromolecules, including nucleic acids, proteins, and phospholipids^4^. Beyond simple charge neutralization, polyamines regulate DNA topology and chromatin architecture^5,6^, modulate transcriptional programs^7,8^, and enhance translational efficiency and fidelity^9^. They participate in numerous other cellular processes, including autophagy, modulation of ion-channel conductance, and maintenance of redox homeostasis^10–13^. The polyamine spermidine provides the exclusive substrate for eukaryotic initiation factor 5A (eIF5A) hypusination: a conserved post-translational modification important for translation elongation and termination^14,15^. Given this breadth of function, cells maintain polyamine abundance under tight homeostatic control.

Cells meet their polyamine requirements through two main strategies: *de novo* synthesis and transport from extracellular sources^2^. Biosynthesis is controlled by a key rate limiting enzyme, ornithine decarboxylase (ODC). ODC has an exceptionally short half-life, allowing for rapid adjustments in polyamine production in response to cellular demands^2^. ODC levels are tightly regulated through a sophisticated feedback loop. Elevated polyamines stimulate antizyme synthesis via programmed ribosomal frameshifting^16^. Antizyme then targets ODC for ubiquitin-independent proteasomal degradation^17,18^. Complementing the biosynthetic pathway, cells acquire extracellular polyamines through specialized transport systems, including P-type ATPases (ATP13A2 and ATP13A3 in mammals, and CATP-5 in nematodes) and other solute carriers^19–23^. The relative contribution of biosynthesis versus import in maintaining polyamine homeostasis across different cell types *in vivo* remains unclear.

Polyamine levels vary markedly with tissue identity and cell state, highlighting the need for tissue-specific regulation. Rapidly proliferating tissues, for instance, demand and maintain constitutively elevated polyamine concentrations: intestinal crypts require high baseline synthesis to support continuous renewal, and regenerating tissues upregulate ODC activity within hours of injury^24–27^. Likewise, CD4+ T cells increase polyamine biosynthesis upon activation, and this surge is essential for rapid clonal expansion and differentiation^8^. Cancer cells exploit these proliferation-polyamine programs to sustain malignant growth^28^. In contrast, differentiated post-mitotic tissues exhibit minimal biosynthetic activity and maintain lower polyamine levels^29,30^

The physiological importance of tissue-specific polyamine regulation is also underscored by focal pathologies arising from systemic disruption. For example, mice lacking spermine synthase (SMS) exhibit hearing loss due to impaired endocochlear potential, a phenotype that mirrors the hearing loss observed in humans treated with ODC inhibitor α-difluoromethylornithine (DFMO)^31^. Likewise, mutations in the broadly expressed polyamine transporter gene, *ATP13A2*, manifest specifically as neurodegenerative disorders including Kufor-Rakeb syndrome and juvenile-onset amyotrophic lateral sclerosis^32–36^. The selective vulnerability of certain tissues to polyamine dysregulation, despite ubiquitous expression of polyamine machinery, suggests that different cell types have distinct polyamine requirements and likely employ tissue-specific regulatory paradigms. Yet we lack basic information about how cell-specific polyamine states are established, maintained, and regulated. Unraveling this regulatory logic is essential not only for understanding normal tissue physiology but also for addressing the selective tissue vulnerabilities observed in polyamine-related diseases.

This current knowledge gap stems at least in part from technical constraints: prevailing methods for measuring polyamines, such as by chromatography or by mass spectrometry, require substantial sample quantities and necessitate tissue destruction^37,38^. These approaches average out the metabolic contributions of individual cells and cannot capture dynamic temporal changes. To address these limitations, we and others have recently developed genetically encoded polyamine sensors that enable real-time, single-cell resolution measurements in live cells^39,40^. In brief, we utilized the polyamine-sensitive frameshift element from the mammalian OAZ1 gene to develop a ratiometric fluorescence reporter, enabling live-cell polyamine measurements at single-cell resolution in cultured mammalian cells^40^.

Here, we adapted our OAZ1 frameshift-based polyamine reporter for use in the nematode *Caenorhabditis elegans*. We generated an *in vivo*, single-cell resolution map of polyamine levels in intact animals and uncovered striking tissue-specific regulation of polyamine homeostasis. By combining this reporter with systematic genetic manipulations, we discovered that most differentiated tissues rely primarily on polyamine transporters rather than biosynthesis to meet their polyamine requirements, with different cell types showing distinct transporter dependencies. Notably, we identified that the intestine, the site of dietary polyamine absorption, serves as the central hub that controls systemic polyamine homeostasis. Finally, longitudinal single-cell resolution measurements allowed us to dissect heterogeneity among neurons, revealing a developmental rewiring where distinct neuron subtypes lose dependency on a polyamine transporter at specific developmental stages. These findings establish fundamental principles of tissue-specific polyamine regulation and provide a framework for understanding how polyamine metabolism contributes to cell-type specific functions in development, aging, and disease.

## Results

### Genetically encoded polyamine sensor for *C. elegans*

We previously developed a genetically encoded polyamine sensor for mammalian cells by harnessing an endogenous polyamine sensing mechanism in which the level of intracellular free polyamines determines the extent of +1 ribosomal frameshifting in the OAZ1 mRNA (Figure 1A)^40^. We reasoned that adapting this approach can yield a *C. elegans* specific polyamine sensor because: 1) the OAZ1 frameshift mechanism and its titration with polyamine abundance is conserved from fission yeast to humans^41^; and 2) *in vitro* translation assays demonstrate that the *C. elegans oaz-1* fragment exhibits spermidine-responsive +1 frameshifting, providing a molecular foundation for sensor development^41^.

**Figure 1.**
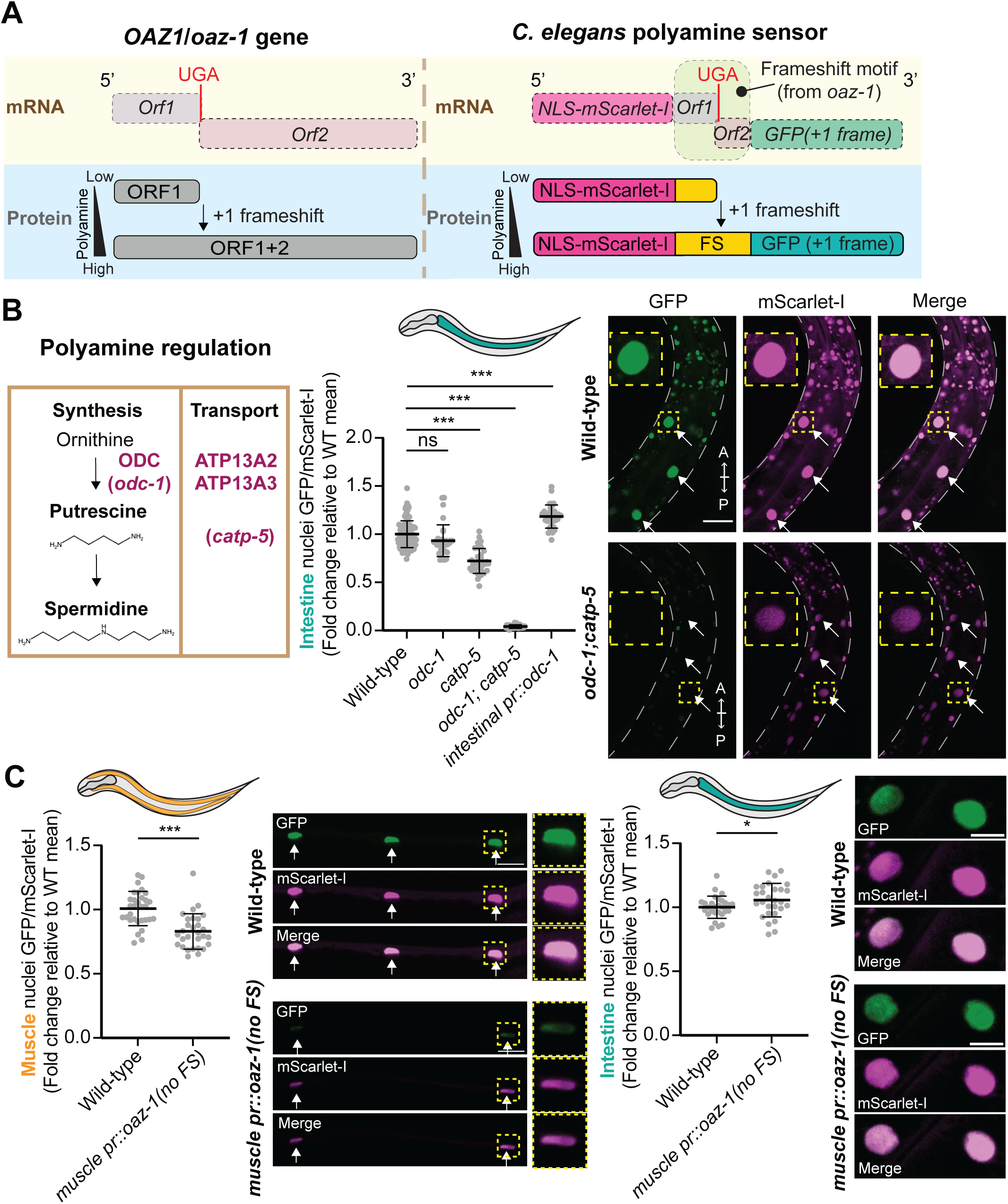
Design and validation of a polyamine sensor in *C. elegans*. (A) Schematic of the polyamine sensor design. (Left) The *oaz-1* gene contains a premature stop codon (in red, UGA) and undergoes a polyamine-dependent +1 ribosomal frameshift to produce full-length OAZ-1 protein (ORF1+ORF2). (Right) Schematic of *C. elegans* polyamine sensor. NLS-mScarlet-I and GFP are appended to either side of the *oaz-1* frameshift motif. Nuclear GFP/mScarlet-I fluorescence ratio reflects intracellular polyamine levels. NLS, nuclear-localization signal. FS, frameshift motif, ORF/*Orf*, open reading frame. (B) (Left) Polyamine synthesis and transport components in human and *C. elegans*. (Middle) Quantification of nuclear GFP/mScarlet-I in intestinal cells of animals with the indicated genotypes. Values are normalized to the mean value of wild-type animals. *C. elegans* cartoon indicates the analyzed tissue. (Right) Fluorescence images of the intestine expressing polyamine sensor in WT and *odc-1; catp-5* animals. Yellow arrow and inset highlight intestinal nuclei. Scale bar, 25 μm. A, anterior direction (of animal); P, posterior direction (of animal). (C) Sensor captures tissue-specific changes in polyamine levels. Quantification of nuclear GFP/mScarlet-I ratio with representative fluorescence images in body wall muscle (Left) or intestine (Right) in WT vs. animals expressing *oaz-1* lacking the frameshift nucleotide under a muscle-specific promoter (p*myo-3*). Values are normalized to the mean value of wild-type animals. Scale bar, 10 μm. (B,C) Each data point (n) is an average of nuclear GFP/mScarlet-I of the tissue of interest in one animal. n= 28-94 across at least three biological replicates. Error bars denote mean ± standard deviation (SD). ns, not significant; *, p≤ 0.05; ***, p≤0.001 ((B) Kruskal-Wallis test with Dunn’s test for multiple comparisons. (C) Mann-Whitney test)). WT, wild-type.

To implement this sensor, we selected a *C. elegans oaz-1* fragment that preserves the polyamine-responsive +1 ribosomal frameshift, spanning 123 bp upstream and 107 bp downstream of the frameshift site (see Materials and Methods, Figure S1A). We deliberately excluded the *oaz-1* C-terminus that binds ODC1 and regulates its activity and degradation (Figure S1A)^42,43^. The reporter was constructed by flanking this *oaz-1* fragment with fluorescent proteins. Upstream, we incorporated a nuclear localization sequence (NLS) and mScarlet-I coding sequence. Downstream, we inserted the GFP coding sequence (without its AUG start codon) into the +1 reading frame (Figure 1A). Ribosomes translating mScarlet-I would encounter an in-frame stop codon. GFP production necessitates a +1 ribosomal frameshift, the efficiency of which increases with intracellular polyamine levels. This design creates a ratiometric reporter where the GFP/mScarlet-I fluorescence ratio reflects polyamine status. The nuclear localization simplifies single-cell quantification and the fusion architecture ensures that GFP signal derives from *bona fide* frameshift-dependent fusion protein expression.

We first expressed the *C. elegans* polyamine sensor under the ubiquitous *eft-3* promoter as a single-copy insertion^44^. mScarlet-I and GFP signals localized to the nucleus, as expected (Figure 1B). Sensor expression did not impact gross animal physiology, including brood size or developmental timing (Figure S1B). To test that the ratiometric readout responds to cellular polyamine status, we leveraged genetic mutants with characterized polyamine deficiencies. *C. elegans* maintains polyamine homeostasis through both biosynthesis and transport (Figure 1B). Abrogation of polyamine synthesis by knockout of polyamine synthesis enzyme *odc-1* can be compensated for by increased polyamine uptake through the CATP-5 transporter, which is localized to the apical side of the gut lumen (Figure 1B)^22,45^. *odc-1; catp-5* double mutants with compromised polyamine synthesis and uptake have substantially reduced polyamine levels compared to either single mutant and exhibit severe developmental delay, body size, and lifespan defects^22^.

We expressed and measured the sensor’s ratiometric readout (GFP/mScarlet-I fluorescence ratio) in intestinal cells of wild-type, *odc-1* and *catp-5* single mutants, and *odc-1; catp-5* double mutant animals. Compared to wild-type, the sensor detected a modest but statistically significant drop in polyamine levels in *catp-5* single mutants, but not in *odc-1* mutants (Figure 1B). Strikingly, the sensor signal in the *odc-1; catp-5* double mutants plummeted to a baseline level (Figure 1B), indicating that the sensor can capture modest changes in polyamine levels in single mutants, and recapitulates the severe, synergistic deficiency that results from disrupting both biosynthesis and import.

Next, we tested if the sensor could detect tissue-specific polyamine changes. To increase polyamines, we overexpressed the biosynthetic enzyme *odc-1* specifically in the intestine, which resulted in a corresponding increase in the sensor ratio in the intestine (Figure 1B). Conversely, to decrease polyamines, we overexpressed a constitutively active form of OAZ-1 (lacking the frameshift site) in body wall muscle. This led to decreased levels of polyamines specifically in muscle, consistent with OAZ-1’s established role in inhibiting polyamine synthesis and import, but had minimal effect on the intestine (Figure 1C)^46^. Together, these results show that our sensor provides a tissue-specific, single-cell readout of intracellular polyamine levels *in vivo*.

Notably, the NLS-mScarlet-I signal, which we originally designed to be a stable expression control, was not constant and instead correlated with polyamine levels (Figures S1C, S1D). We suspect that this is due to proteasomal degradation of the NLS-mScarlet-I protein when it is not extended by the GFP-containing +1 frame product (Figure S1E) (see Supplementary note). This regulatory degradation of the nascent OAZ1 frame-0 peptide was recently reported in yeast and mouse cells^47^. This behavior introduces a potential confounder: under low-polyamine conditions, reduced mScarlet-I would artificially elevate the GFP/mScarlet-I ratio and could, in principle, mask a decrease in frameshifting. Nonetheless, we observed a significant decrease in the GFP/mScarlet-I ratio under low-polyamine conditions (Figure 1B), indicating that the reduction in frameshift-dependent GFP expression is substantial enough to overcome the confounding effects from mScarlet-I changes. These data confirm that the ratiometric output, considered alongside the direction of mScarlet-I changes, is a reliable indicator of intracellular polyamine status. We provide both the GFP/mScarlet-I ratios and the mScarlet-I fluorescence measurements for all experiments in the supplementary data.

### Somatic tissue polyamine measurements reveal low polyamine levels in neurons

Having established that our polyamine sensor reports on tissue-specific polyamine levels, we next leveraged it to examine whether different cell types maintain distinct polyamine setpoints *in vivo*. We expressed our polyamine sensor using tissue-specific promoters in four major somatic tissues: neurons, body wall muscle, intestine, and the syncytial hypodermis. To ensure we were measuring homeostatic levels in stable cell populations, we conducted our analysis at the larval stage 4 (L4), just prior to adulthood, when somatic cell divisions have ceased in most tissues^48^. Sensor measurements revealed that polyamine levels were similar in the intestine, body-wall muscle, and hypodermis, but markedly lower in neurons (Figure 2B). To distinguish genuine differences in polyamine-dependent frameshifting from potential tissue-specific variations in promoter activity or translation efficiency, we engineered a control reporter expressing an mScarlet-I-GFP fusion protein without the intervening frameshift site. This control exhibited nearly uniform fluorescence ratios across all tissues examined, confirming that our sensor measurements reflect true differences in cellular polyamine content rather than differences in transcriptional activity or cellular translation rates (Figure 2C). This finding demonstrates that even among post-mitotic, non-dividing cell types, polyamine content can vary significantly. Overall, these results show that polyamine homeostasis is not uniformly maintained across differentiated tissues but is subject to tissue-specific regulation, likely reflecting their distinct functional and metabolic demands.

**Figure 2.**
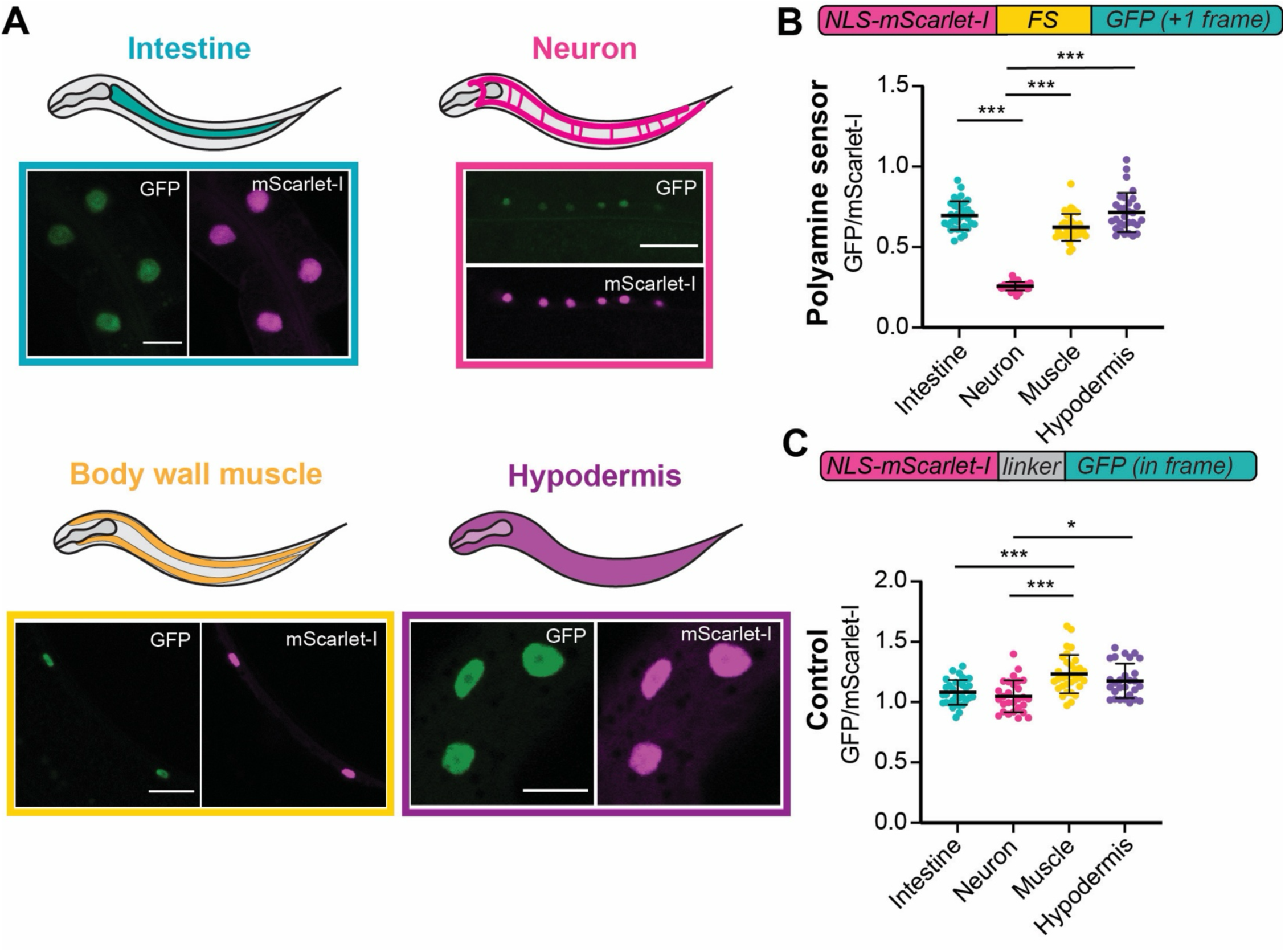
Neurons exhibit lower polyamine levels compared to other somatic tissues. (A) Visualization of polyamine sensor readout in various somatic tissues. Representative fluorescence images show sensor output in intestine, neurons, body wall muscle, and hypodermis at the L4 stage. Tissue-specific expression was driven by the following promoters: intestine, p*vha-6*; neuron, p*unc-11prom8*; muscle, p*myo-3*; and hypodermis, p*col-10*. Scale bar, 10 μm. (B,C) Quantification of nuclear GFP/mScarlet-I fluorescence ratios in the indicated tissue types at the L4 stage using the polyamine sensor (B) or control construct (C). n= 28-34 animals per genotype across three biological replicates. Only statistically significant comparisons are shown. *, p≤ 0.05; ***, p≤0.001 (Kruskal-Wallis test with Dunn’s test for multiple comparisons). Error bars denote mean ± SD. FS, frameshift motif.

### Polyamine transport is the major pathway used for polyamine maintenance in somatic tissues

The heterogeneity in polyamine levels between tissues prompted us to ask whether different tissues use distinct mechanisms to establish and maintain their polyamine setpoints. Surprisingly, similar to our observation in the intestine (Figures 1B), knockout of *odc-1*, encoding the primary biosynthetic enzyme, did not substantially alter polyamine levels in muscle, neurons, or hypodermis (Figure 3A, S2A). This result suggested that either transport can effectively compensate for reduced *de novo* synthesis or it might be the dominant polyamine acquisition route in these somatic tissues.

**Figure 3.**
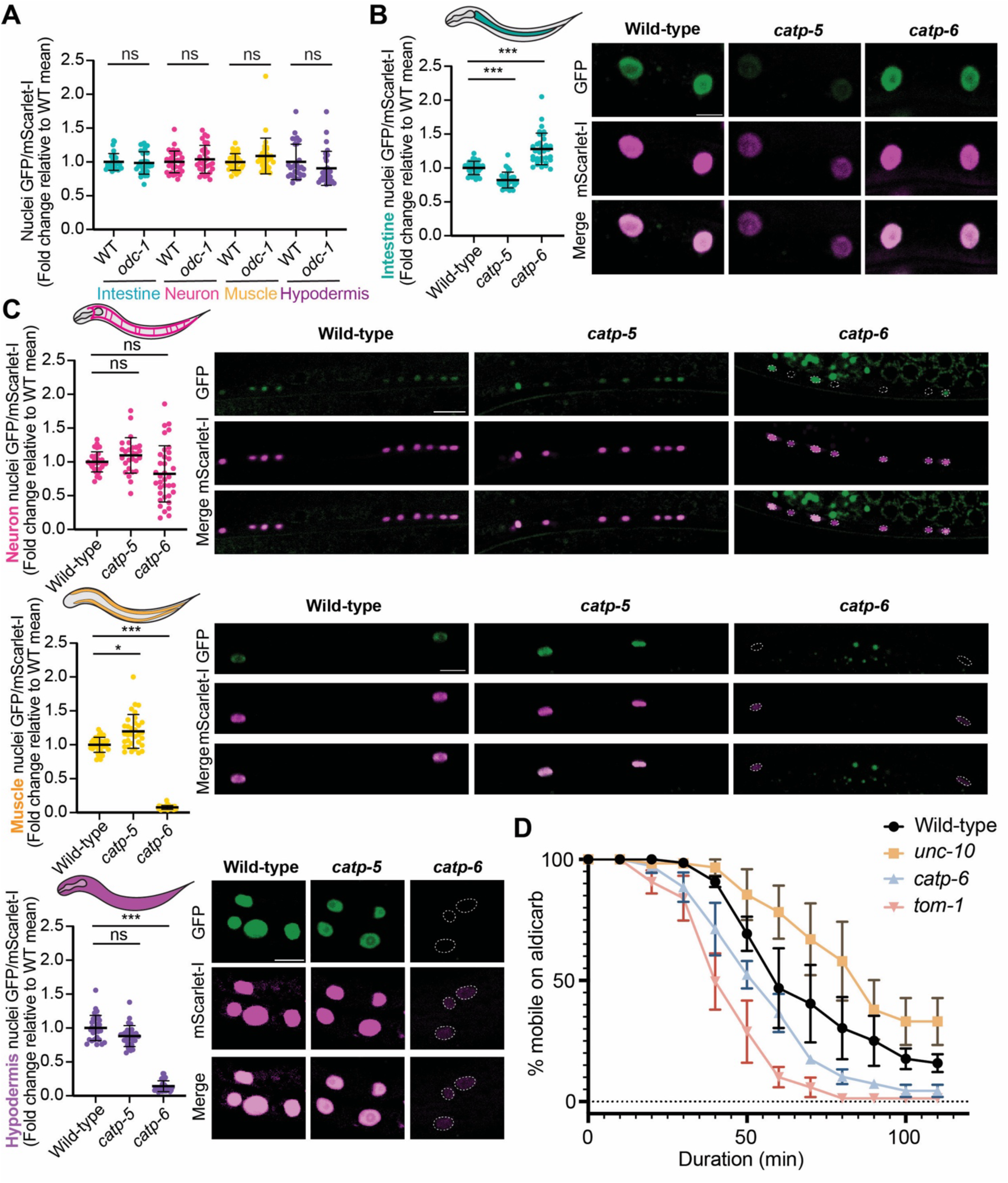
Polyamine transporters are the primary modulators of polyamine levels in somatic tissues. (A) Quantification of polyamine levels in indicated tissues in wild-type and *odc-1* mutants. Values are normalized to the mean value of wild-type animals. (B) (Left) Quantification of intestinal polyamine levels in wild-type and polyamine transporter mutants. Values are normalized to the mean value of wild-type animals. (Right) Representative fluorescence images show corresponding sensor expression in intestinal nuclei. Scale bar, 10 μm. (C) Tissue-specific impact of polyamine transporter mutations on sensor output. (Left) Quantification of neuron, muscle, and hypodermis nuclear GFP/mScarlet-I in wild-type and polyamine transporter mutants expressing polyamine sensor. Values are normalized to the mean value of wild-type animals. (Right) Representative fluorescence images of neuron, muscle, hypodermis nuclei of wild-type and polyamine transporter mutants expressing polyamine sensor. Scale bar, 10 μm. (A-C) n=27-36 animals per genotype across three biological replicates. ns, not significant; *, p≤ 0.05; ***, p≤0.001 (Kruskal-Wallis test with Dunn’s test for multiple comparisons.) Error bars denote mean ± SD. (D) Aldicarb-induced paralysis assay of WT vs. *catp-6* mutants. *unc-10* and *tom-1* mutants are controls for resistance and hypersensitivity to aldicarb respectively. 18-25 animals per genotype scored per round, with a total of three rounds scored on separate days. Error bars denote mean ± standard error of the mean.

In *C. elegans*, there are three orthologs of the mammalian polyamine transporters ATP13A2 and ATP13A3: P5B-type ATPases *catp-5*, *catp-6*, and *catp-7*^22,49^. To determine the tissue-specific transporter dependencies, we used CRISPR/Cas9 to create putative null alleles of each transporter and measured polyamine levels across tissues using our sensor. This analysis revealed prominent tissue selectivity in transporter usage. Consistent with the predominantly intestinal expressions of *catp-5* mRNA and CATP-5 protein^49–52^, *catp-5* knock out modestly decreased polyamine levels in the intestine but did not affect neurons, body wall muscle, or hypodermis (Figures 3B, 3C, S2B, S2C). In contrast, *catp-6* knock out dramatically reduced polyamines in body wall muscle, hypodermis, and a subset of neurons (characterized in detail below) (Figures 3C, S2C), consistent with a broad expression of CATP-6 (Figure S2B)^53^. Intestinal polyamine levels were elevated in *catp-6* mutants, possibly reflecting compensatory upregulation in response to systemic polyamine deficiency (Figures 3B, S2C). *catp-7* knock out did not affect neurons or intestine (Figure S2D), but resulted in a modest reduction in body wall muscle polyamine levels (Figure S2D).

The severe polyamine depletion in muscle and neurons upon loss of *catp-6* is in stark contrast to the normal levels maintained in *odc-1* mutants and suggests that these tissues are exceptionally reliant on transport to maintain polyamine homeostasis. To test if this molecular deficiency has physiological consequences, we assessed neuromuscular function using aldicarb-induced paralysis assays. Aldicarb is an acetylcholinesterase inhibitor that causes acetylcholine to accumulate at the synaptic cleft, leading to muscle hyper-contraction and paralysis^54^. *catp-6* mutants displayed hypersensitivity to aldicarb, with accelerated paralysis compared to wild-type animals (Figure 3D). This phenotype is consistent with synaptic transmission defects and demonstrates that tissue-specific polyamine levels are functionally important^54^. Notably, a previous study also showed that knockdown of *catp-6* increased α-synuclein aggregation in *C. elegans* body wall muscles^55^. Together, these data reveal that transport, not *de novo* synthesis, predominantly sets steady-state polyamine levels in somatic tissues, with *catp-5* and *catp-6* providing distinct, tissue-matched transport routes.

### *C. elegans* AZIN-1 primarily modulates the polyamine transport pathway

We were surprised by the observation that *odc-1* knock out did not affect tissue polyamine homeostasis. While investigating the polyamine biosynthetic machinery, we made an intriguing observation: *C. elegans* ODC-1 lacks the C-terminal degron sequence that confers the characteristic short half-life of mammalian ODC (Figures 4A, S3A)^56–59^. This suggests that *C. elegans* ODC-1 may be more stable than its mammalian counterpart, as is the case for *Trypanosoma brucei* ODC, which also lacks the C-terminal tail (Figure S3A)^57^. This structural difference prompted us to search for additional regulatory mechanisms.

**Figure 4.**
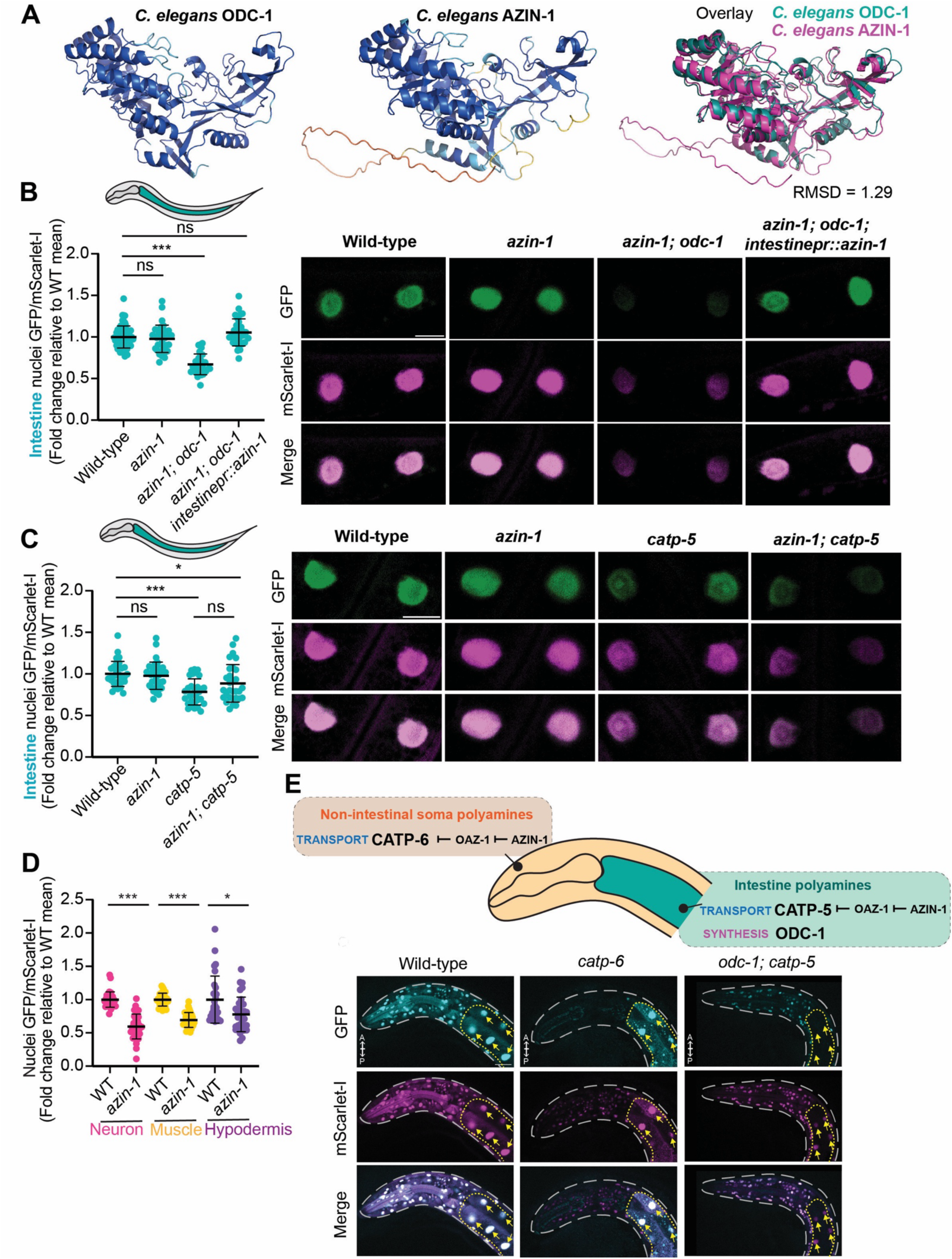
AZIN-1 regulates polyamine transport. (A) Alphafold2 structure predictions of *C. elegans* ODC-1 and AZIN-1^58,59^. Overlay of the models reveals high structural similarity (RMSD, Root Mean Square Deviation = 1.29 Å). Coloring represents predicted local distance difference test (pLDDT) values by Alphafold color scheme (https://github.com/sokrypton/ColabFold). (B,C) (Left) Quantification of intestinal polyamine levels in indicated genotypes. Values are normalized to the mean value of wild-type animals. (Right) Representative fluorescence images show corresponding sensor expression in intestinal nuclei. Scale bar, 10 μm. (D) Quantification of neuron, muscle, and hypodermis polyamine levels in animals of indicated genotypes. Values are normalized to the mean value of wild-type animals. (B-D) n= 29-33 per genotype across at least three biological replicates. ns, not significant; *, p≤ 0.05; ***, p≤0.001 (Kruskal-Wallis test with Dunn’s test for multiple comparisons). Error bars denote mean ± SD. (E) Summary of somatic tissue polyamine regulation: Non-intestinal tissues use *catp-6* mediated transport as their major means of acquiring polyamines; Intestinal tissues use both *catp-5* mediated transport and polyamine synthesis to maintain polyamine levels. Fluorescence images of anterior regions of animals expressing polyamine sensor in all somatic cells. Intestine is outlined in yellow; intestinal nuclei are indicated by yellow arrows. All other nuclei belong to non-intestinal soma. Note that in *catp-6* mutants, polyamine levels are drastically decreased in all non-intestinal somatic tissues. In *odc-1; catp-5* mutants, polyamine levels are decreased most dramatically in the intestine. Scale bar, 20 μm. A, anterior direction; P, posterior direction (of animal).

While searching for additional ODC homologs in the *C. elegans* genome, we found the putative *C. elegans* antizyme inhibitor, *azin-1*, which unlike ODC-1, contains the characteristic C-terminal tail found in mammalian ODC (Figures 4A, S3B)^60^. Antizyme inhibitors are structurally similar to ODC, and typically function by competing for antizyme binding, thereby preventing ODC degradation and maintaining polyamine biosynthetic capacity^61^. Interestingly, in *C. elegans*, single-cell RNA sequencing data show largely non-overlapping expression patterns of *odc-1* and *azin-1* genes: Although both proteins are expressed in the intestine, *azin-1* shows prominent expression in neurons that have relatively low *odc-1* expression (Figure S3C)^50–52^. This expression pattern suggested that AZIN-1 in worms may not principally act by regulating ODC-1.

In mammalian cells, besides inhibiting ODC, antizyme has also been shown to inhibit polyamine transporter activity^61^. Since AZIN-1 sequesters antizyme, it can potentially prevent antizyme-mediated inhibition of ODC as well as polyamine transporters. We therefore hypothesized that *C. elegans* AZIN-1 regulates polyamine transport. To test this notion, we performed genetic epistasis analysis in the intestine, asking whether *azin-1* functions in synthesis (*odc-1*) or transport (*catp-5*) pathways. Similar to what we observed in *odc-1* single mutants, *azin-1* knock out did not measurably alter polyamine levels in the intestine (Figures 4B, S3D). In contrast, *azin-1; odc-1* double mutants exhibited polyamine depletion compared to the single mutants alone (Figures 4B, S3D). Crucially, *azin-1; catp-5* double mutants showed no additive effects compared to *catp-5* single mutants, suggesting that AZIN-1 likely functions in the transport pathway (Figures 4C, S3E).

The physiological importance of AZIN-1’s transport regulatory function became apparent when we examined non-intestinal somatic tissues that rely heavily on polyamine transporters. While *odc-1* mutants maintained normal polyamine levels in neurons, muscle, and hypodermis, *azin-1* mutants showed reduced polyamine levels in all three tissues (Figures 4D, S3F, S3G). These data collectively reveal a new regulatory logic: in *C. elegans*, *azin-1* likely exerts its effect primarily through modulating polyamine transporter function, rather than its canonical mechanism of protecting ODC from OAZ-mediated degradation. These data are also consistent with our model that while the intestine maintains functional redundancy between synthesis and transport pathways, other tissues such as neurons, muscle, and hypodermis rely almost exclusively on transporter-mediated polyamine import (Figure 4E).

### The intestine serves as a central hub for organismal polyamine homeostasis

Given that the intestine uses redundant mechanisms for maintaining intracellular polyamine levels and is the gateway for dietary polyamine absorption, we hypothesized that it might serve as a major hub for polyamine homeostasis in the entire animal. To test this, we re-examined the *odc-1; catp-5* double mutants, where both polyamine synthesis and import are blocked and animals exhibit drastic decreases in intestinal polyamine levels (Figure 1B). We confirmed that the intestinal polyamine depletion in *odc-1; catp-5* double mutants could be rescued by intestine-specific restoration of ODC-1, but not by intestine-specific expression of ODC-like antizyme inhibitor, AZIN-1 (Figures S4A, S4B). Interestingly, *odc-1; catp-5* double mutants displayed a systemic polyamine deficiency that extended beyond the intestine. For example, polyamine levels were significantly reduced in the head region, which consists of various somatic cell types including neurons, muscle, and hypodermis (Figures 5A, S4C). This was particularly surprising given that *odc-1* knockout alone does not reduce polyamine levels in these tissues (Figure 3A), and that these tissues primarily import polyamines via CATP-6, not CATP-5 (Figure 3C). This systemic deficiency may explain at least in part the prominent phenotypes in *odc-1; catp-5* double mutants, including decreased body size, diminished brood size, and delayed development (Figure 5B)^22^.

**Figure 5.**
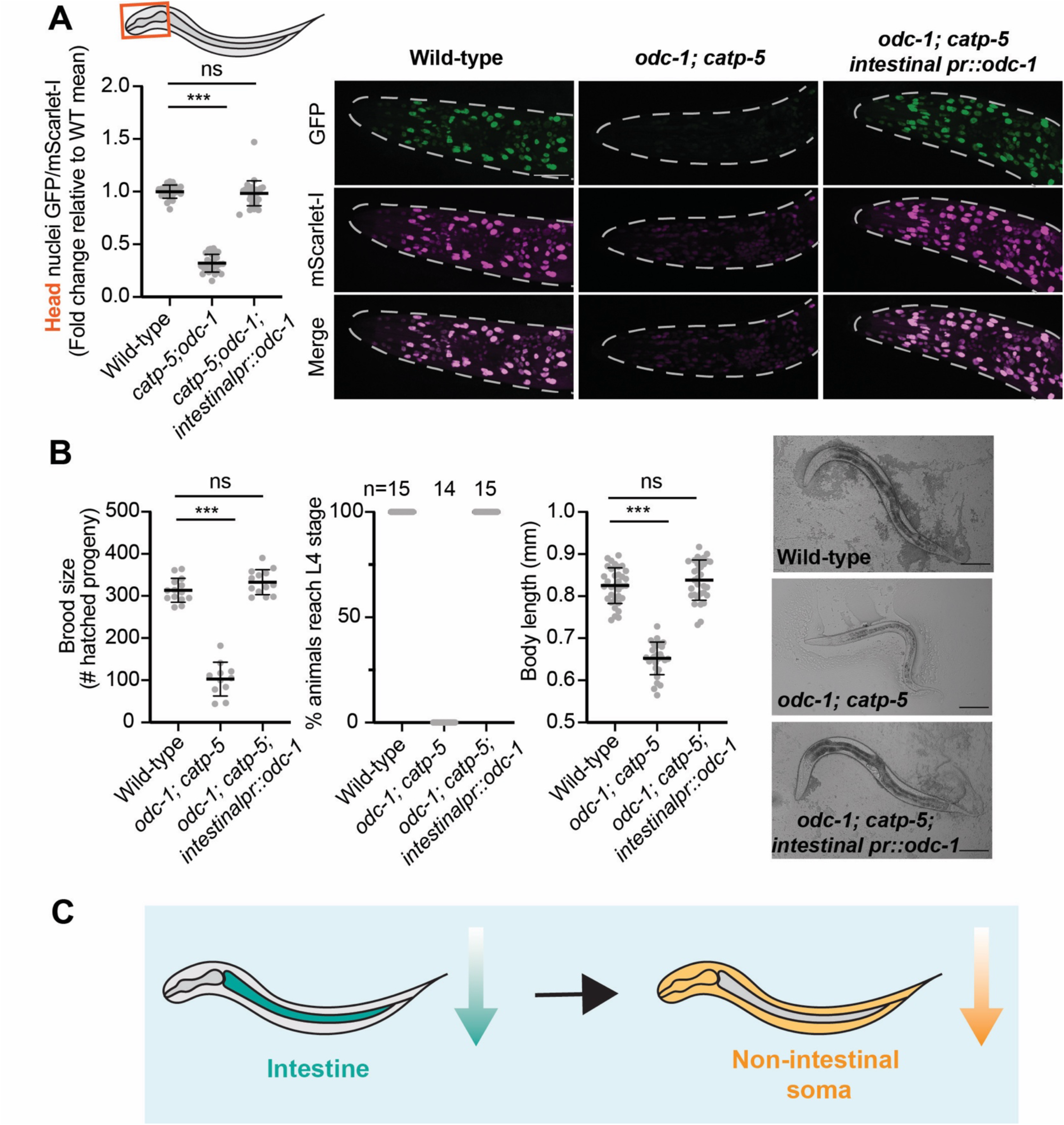
The intestine is a central hub for organismal polyamine regulation. (A) (Left) Quantification of polyamine levels in head cells of indicated genotypes. Values are normalized to the mean value of wild-type animals. n=25-29 per genotype across three biological replicates. (Right) Representative fluorescence images show corresponding sensor expression in head nuclei. Scale bar, 25 μm. (B) Brood size (n=10 per genotype across three biological replicates) (left graph), developmental timing (middle graph), and body length (n=30-34 per genotype across three biological replicates) (right graph) of indicated genotypes. Development timing was scored as percentage of progeny animals reaching L4 stage after ∼52 hours post egg stage. n indicates sample size (14-15 plates per genotype, with at least 11 progeny animals scored per plate). (Right) Images of animals to show body length. Scale bar, 0.1 mm. (A,B) ns, not significant; ***, p≤0.001 (Kruskal-Wallis test with Dunn’s test for multiple comparisons). Error bars denote mean ± SD. (C) Summary of inter-tissue polyamine regulation: When intestinal polyamine levels are lowered, non-intestinal soma polyamine levels are decreased.

Using tissue-specific rescue experiments, we found that restoring *odc-1* expression exclusively in the intestine was sufficient to restore polyamine balance throughout the entire animal as well as rescue all systemic phenotypes (Figures 5A, 5B, S4C). These results demonstrate that intestinal polyamine reserves are not just used locally, but support polyamine levels systemically, either through direct transport or other indirect mechanisms. The fact that non-intestinal tissues cannot maintain normal polyamine levels when intestinal polyamines are depleted, despite having intact transport systems, highlights the importance of intestinal polyamine reserves for organismal health (Figure 5C). More broadly, our ability to visualize this inter-tissue metabolic regulation highlights how genetically encoded sensors applied *in vivo* can reveal systemic regulatory architectures that would be invisible in bulk measurements or *in vitro* applications.

### Developmental remodeling of neuronal polyamine transport dependencies

A major advantage of genetically encoded sensors over traditional biochemical approaches is the ability to track metabolic dynamics within the same cells over time. Our analyses thus far have captured a static snapshot of polyamine levels at a single developmental timepoint, the L4 stage. We next harnessed this longitudinal tracking capability to examine whether polyamine regulation undergoes developmental changes.

We chose three time points to assess polyamine levels: 1) early larval stage (24 hours after the egg was laid); 2) L4 stage; 3) young adult stage (24 hours post L4 stage) (Figure 6A). Anatomically, we focused on the ventral nerve cord (VNC). The VNC has been likened to the *C. elegans* equivalent of the vertebrate spinal cord, and comprises two classes of motor neurons: cholinergic (acetylcholine-releasing) or GABAergic (GABA-releasing), with the exception of VC4 and VC5^48^ (Figure 6A). GABAergic neurons can be identified using a neuronal marker (p*unc-25*::NLS::mTagBFP2), which is not expressed in non-GABAergic neurons (including cholinergic neurons, VC4, and VC5).

**Figure 6.**
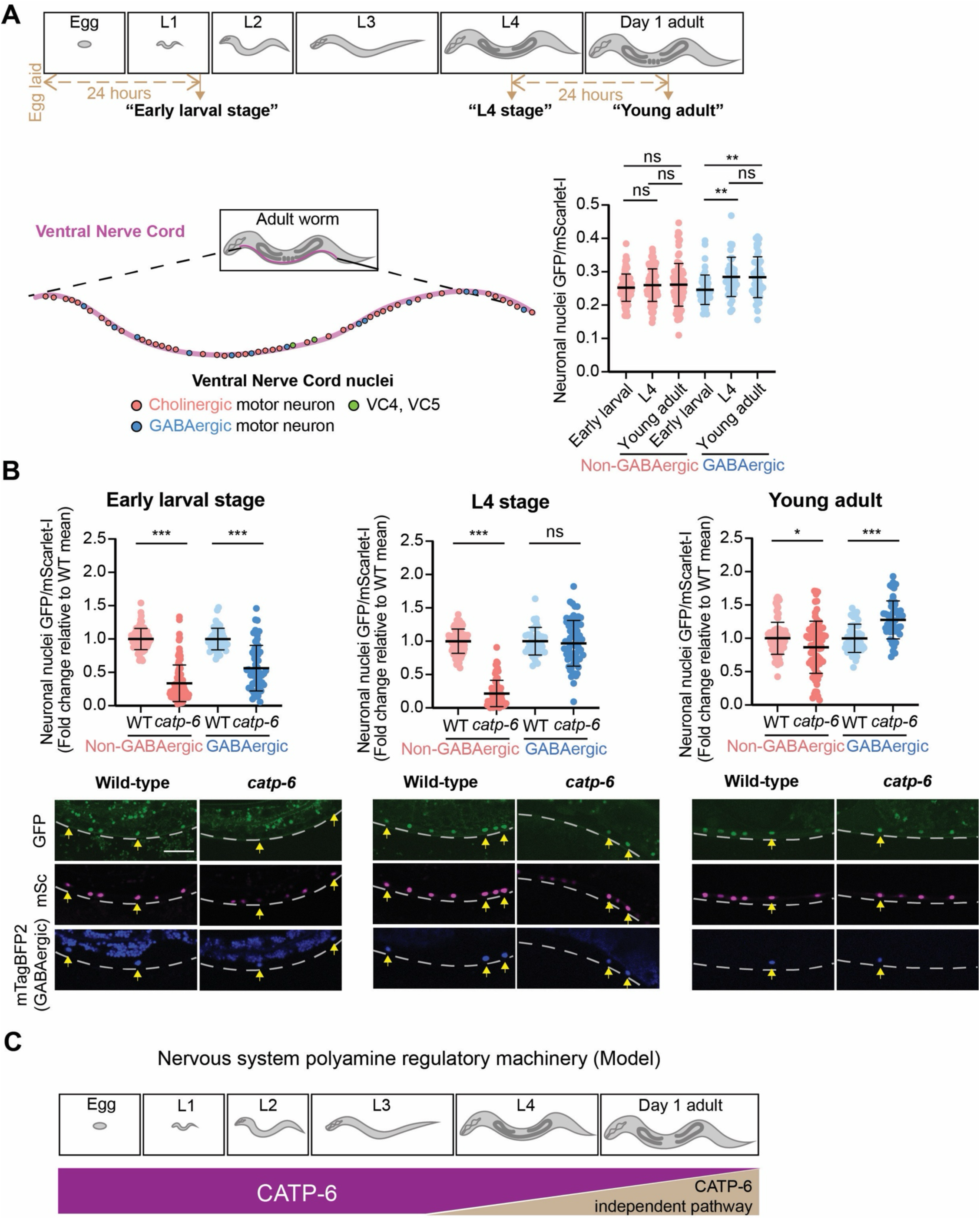
Polyamine transporter dependency switches during nervous system development. (A) (Top) Schematic of developmental time-points assayed. *C. elegans* goes through four larval stages before adulthood. “Early larval stage” refers to 24 hours post egg stage; “Young adult stage” refers to 24 hours post L4 stage. (Bottom left) Schematic of anatomy of ventral nerve cord (VNC) nuclei: VNC consists purely of motor neuron nuclei, including cholinergic (acetylcholine-releasing) neurons, GABAergic (GABA-releasing) neurons, and serotonin-releasing VC4, VC5. (Bottom right) Quantification of polyamine levels in VNC non-GABAergic and GABAergic neurons across developmental time points. (B) (Top) Quantification of polyamine levels in VNC non-GABAergic and GABAergic neurons in wild-type and *catp-6* mutants across developmental time points. Values are normalized to the mean value of the same neuron type and of the same developmental stage in wild-type animals. (Bottom) Representative fluorescence images show corresponding sensor expression in VNC nuclei. GABAergic neuronal nuclei are marked with mTagBFP2 and indicated with yellow arrows. Scale bar, 15 μm. (A,B) Each data point (n) is a neuronal nuclei GFP/mScarlet-I value. n= 40-94 from 30-35 animals for each condition (genotype/stage/neuron type combination) across at least three biological replicates. Error bars denote mean ± SD. ns, not significant; *, p≤ 0.05; **, p≤ 0.01; ***, p≤0.001 ((A) Kruskal-Wallis test with Dunn’s test for multiple comparisons. (B) Mann-Whitney test). (C) Model of developmental switching of polyamine regulatory program: At stages prior to L3/L4, the nervous system mainly uses CATP-6 to maintain polyamine levels. This reliance decreases as animals enter adulthood.

Tracking these defined neuronal populations across development revealed dynamic changes in polyamine homeostasis. We observed a small but significant increase in polyamine levels in GABAergic motor neurons from early larval to L4 stage, while non-GABAergic neurons maintained relatively stable levels (Figure 6A). This selective change in basal polyamine levels in GABAergic neurons suggested potential developmental remodeling of polyamine metabolism.

Interestingly, during our initial analysis of *catp-6* mutants (Figure 3C), we observed substantial heterogeneity in polyamine levels specifically within the VNC. We hypothesized that this variability reflects neuron subtype-specific differences in *catp-6* dependence. Indeed, temporal analysis revealed a developmental program: at early larval stage, *catp-6* loss decreased polyamine level in both non-GABAergic and GABAergic motor neurons compared to wild-type animals, indicating an initial dependence on *catp-6* in both neuron types (Figures 6B, S5A). However, at the L4 stage, a divergence emerged: *catp-6* was no longer required for polyamine level maintenance in GABAergic motor neurons, while non-GABAergic neurons remained severely polyamine-depleted (Figures 6B, S5A). This differential response was not due to differences in sensor expression or translation as control experiments in *catp-6* mutant animals using a frameshift independent NLS-mScarlet-I-GFP fusion protein construct showed comparable nuclear GFP/mScarlet-I ratios between non-GABAergic and GABAergic motor neurons at L4 stage (Figure S5B).

Most remarkably, this metabolic reprogramming continued into adulthood. At the young adult stage, non-GABAergic motor neurons also showed the trend of achieving *catp-6* independence (Figures 6B, S5A). Thus, polyamine regulatory mechanisms undergo significant developmental remodeling in the nervous system, with different neuron types following distinct temporal trajectories in their reliance on specific transport pathways (Figure 6C). While we did not quantify polyamine levels for other neuron types, qualitatively, neurons in the head and tail regions of *catp-6* mutant animals followed similar developmental trajectory as non-GABAergic motor neurons (Figures S5C-S5E), suggesting that the developmental progression is not unique to VNC motor neurons but represents a nervous system-wide phenomenon. Together, our findings reveal a previously unappreciated complexity in polyamine regulation that varies with both cell type and developmental stage, highlighting the power of our genetically encoded reporter and single-cell measurements in understanding polyamine biology in multicellular organisms.

## Discussion

We developed and deployed a genetically encoded polyamine sensor in *C. elegans* to generate a single-cell resolution map of polyamine levels in a living multicellular organism. This technological advance overcomes longstanding limitations of bulk measurement techniques that require removal of cells from their native environments and lack the ability to capture spatial heterogeneity or temporal dynamics^37,38^.

We find that different tissues maintain distinct polyamine setpoints. Neurons exhibit substantially lower polyamine levels compared to other differentiated tissues including muscle, intestine, and hypodermis. This heterogeneity challenges the prevailing view that polyamine requirements primarily correlate with proliferative status. This observation may provide a metabolic basis for the selective vulnerability of the nervous system to polyamine dysregulation. Pathologies arising from mutations in ubiquitously expressed polyamine-related genes, such as the transporter ATP13A2 or the catabolic enzyme SSAT, often manifest specifically as neurodegenerative disorders like Parkinson’s disease and amyotrophic lateral sclerosis^32–34,62^. The lower basal polyamine pool may provide a smaller buffer against metabolic changes and even modest disruptions in polyamine levels could produce severe functional consequences in neurons compared to tissues with larger polyamine reserves.

A surprising finding from our study is that most differentiated tissues rely almost exclusively on polyamine transport rather than *de novo* synthesis. Loss of *odc-1*, the rate-limiting biosynthetic enzyme, alone failed to reduce polyamine levels in any of the tissues examined. On the other hand, knock out of specific transporters resulted in modest polyamine depletion, with physiological consequences including synaptic dysfunction. This dependence on transport represents a fundamental shift in how we conceptualize polyamine homeostasis in multicellular organisms, from a focus on cell-autonomous synthesis to tissue programs that tune uptake. Several specialized mammalian polyamine transporters have been identified^19–21,23,63^. Given our observations in *C. elegans*, it is possible that transport regulation, not biosynthesis, plays the key role in polyamine regulation and its diversification amongst mammalian cell types.

Our findings establish the intestine as the central hub for organismal polyamine homeostasis. The intestine is unique in using redundant polyamine synthesis and transport pathways for polyamine maintenance, while other somatic tissues we surveyed rely primarily on polyamine transport. Depleting intestinal polyamines decreased polyamines in non-intestinal somatic tissues and had broad physiological impacts. Crucially, restoring *odc-1* expression exclusively in the intestine was sufficient to rescue polyamine levels throughout the entire animal and correct systemic defects. These results suggest that therapeutic interventions targeting intestinal polyamine metabolism could modulate polyamine levels across tissues. More broadly, these findings underscore the importance of studying metabolism in intact living organisms, where inter-tissue regulation could be identified.

Beyond identifying tissue-specific acquisition strategies, our work revealed unexpected regulatory mechanisms. We found that *azin-1* knock out primarily impacted polyamine transport but not synthesis. While AZIN-1 is best studied for promoting polyamine synthesis by sequestering antizyme (OAZ) and protecting ODC from degradation^64^, OAZ has also been shown to inhibit polyamine transport^65–68^. This raises the possibility that AZIN-1 modulates polyamine transport through the same OAZ-sequestration mechanism. Interestingly, a genome-wide CRISPR-Cas9 knockout screen in mammalian cells for regulators of spermidine uptake identified AZIN1 among the top hits, indicating that AZIN’s function as a polyamine transport modulator is conserved in mammals^40^. Although the precise molecular mechanisms by which OAZ and AZIN regulate polyamine import remain elusive, understanding this interplay has critical therapeutic implications, as AZIN overexpression is observed in tumors and can drive cell transformation^68,69^.

In addition to detecting spatial heterogeneity, our live-animal sensor revealed temporal dynamics in polyamine regulation. We uncovered a developmental switch in neuronal transporter dependencies, right around the time when the nervous system transitions from immature to mature states. This developmental reprogramming parallels the observations in mammalian systems, where activity of polyamine synthesis enzymes and expression of polyamine transporters shift during neuronal maturation^70,71^. For instance, ATP13A2 levels vary across development, with highest expression during neurogenesis^71^. Intriguingly, environmental factors including maternal care quality can also influence offspring polyamine biosynthesis activity^72,73^, suggesting that both genetic programs and early life experiences influence polyamine balance. Understanding the triggers and functional significance of these developmental transitions could reveal critical periods for intervention and guide therapeutic timing for polyamine-related diseases. By directly linking cellular polyamine status to organismal physiology, our sensor may illuminate disease mechanisms, provide an *in vivo* platform for designing and evaluating therapeutic interventions, and may enable us to harness the polyamine pathway to its full pharmacological potential.

### Limitations of the study

While our polyamine sensor provides unprecedented resolution for tracking polyamine dynamics *in vivo*, an important limitation is that it measures relative rather than absolute polyamine concentrations. Converting fluorescence ratios to absolute polyamine levels would require extensive calibration against biochemical measurements. Although we successfully calibrated the sensor in mammalian cell culture^40^, similar calibrations in *C. elegans* would require prohibitively large quantities of sorted cells or dissected tissues to obtain sufficient material for quantitative analysis by chromatography or mass spectrometry. Additionally, the sensor specifically detects polyamines that modulate OAZ1 frameshifting (spermidine and possibly to a lesser extent, putrescine). Despite these constraints, the ability to measure relative changes with single-cell resolution allowed us to gain significant insights into polyamine regulation *in vivo*. As mass spectrometry and related detection methods continue to improve, combining relative measurements from genetic sensors with targeted absolute quantification will provide a comprehensive view of polyamine regulation.

## Materials and Methods

### Strains

*C. elegans* strains were cultivated at 20°C under standard growth conditions on nematode growth medium (NGM) plates with OP50 *Escherichia coli* as food. Single-copy gene insertions were generated by Minimos insertion technique^44^. Polyamine synthesis and transporter mutants were generated by CRISPR/Cas9 technology with polyamine sensor strains as starting strains^74^. CRISPR/Cas9 modifications for the gene knockouts are as follows: For *odc-1* knockout, an 8-base pair (‘taatctag’) sequence was inserted in *odc-1* exon 1, which is upstream of the ODC-1 catalytic active site; for *azin-1* knockout, a 2651 bp deletion encompassing coding sequences of all *azin-1* isoforms was created; for *catp-5* knockout, an 8-base pair (‘taactagt’) sequence was inserted in exon 1 of *catp-5* isoform a, which is upstream of the P-type ATPase actuator (A), phosphorylation (P), nucleotide-binding (N) domains and pore-forming transmembrane domains; for *catp-6* knockout, an 8-base pair (‘taactagt’ or ‘taatctag’) sequence was inserted in exon 2 of *catp-6* isoform b, which is upstream of the P-type ATPase P, N domains, and M4-M10 of the transmembrane domains; for *catp-7* knockout, an 8-base pair (‘taactagt’) sequence was inserted in exon 2 of *catp-7* isoform a, which is upstream of the the P-type ATPase A, P, N domains and pore-forming transmembrane domains^49,75–77^. Guide RNAs were designed using the CRISPOR site^78^. Description of strains used in this study is listed in Supplementary Table 1.

### Polyamine sensor and control plasmid construction

Polyamine sensor plasmid constructs are composed of an ubiquitous or tissue-specific promoter driving a minimal degron sequence (AID*) followed by SV40 nuclear-localization sequence (NLS) and mScarlet-I sequence^79,80^. An 18 bp linker (GGSGGS) and the initial 229 base-pairs of the *oaz-1* coding sequence (excluding start codon) follows, with another 18 bp linker (GGSGGS) downstream linking to a +1 frame GFP sequence (specifically GFP variant S65C) and the *unc-54* 3’UTR. Promoters used are as follows: *eft-3* promoter (622 bp) for expression in all somatic cells, *vha-6* promoter (876 bp) for expression in intestinal cells, *unc-11prom8* promoter (293 bp) for expression in all neurons^81^, *myo-3* promoter (2386 bp) for expression in body wall muscle cells, and *col-10* promoter (640 bp) for expression in hypodermal cells. Control construct plasmid (used in Figures 2C, S5A, Supp Note Fig 1) composes of a tissue-specific promoter driving AID* followed by SV40 NLS::mScarlet-I, a six amino acid linker (encoding the peptide GGSGGS), and in-frame GFP (variant S65C).

### Brood size assay

An animal was singled at L4 stage, and its hatched progeny were counted for the next consecutive 5-8 days. Endpoint of assay was determined by when egg laying ceased. Assays were performed with the researcher blinded to genotype.

### Developmental timing assay

Ten gravid animals were picked onto an NGM plate for egg laying for 3 hours at room temperature (23°C) before removal of the gravid animals. After egg laying, plates were maintained in 20°C for progeny to grow up. ∼52 hours after egg laying, the percentage of progeny animals that reached L4 stage were scored under a dissecting microscope. Assays were scored with the researcher blinded to genotype.

### Animal length measurement

L4 animals were immobilized with polystyrene beads (Polysciences, Cat #00876) and mounted on 10% agarose pads. Images were taken with the UPlanSApo 10x/0.40 objective (Olympus) at brightfield on the EVOS M7000 microscope (Invitrogen). Images were loaded into Adobe Illustrator Version 25.4.1. A line from head to tail of each animal was manually drawn and measured using the measure path length function in Adobe Illustrator.

### Confocal microscopy image acquisition

All polyamine level measurements using the polyamine sensor, except Figures 6 and S5, were performed at the L4 stage. Animals were immobilized with polystyrene beads (Polysciences, Cat #00876) and mounted on 10% agarose pads. Images were taken with the Nikon 60x/1.40 oil immersion objective using the Dragonfly 505 spinning-disk confocal microscope equipped with iXon Ultra 888 EMCCD camera (Andor Technologies). When measuring nuclear GFP/mScarlet-I for comparison between wild-type and mutants in the same tissue, confocal settings (including exposure time, gain, laser intensity) were kept constant between wild-type and mutants for each fluorescence channel used. When measuring nuclear GFP/mScarlet-I for comparison between different tissues of the same genotype (Figure 2), laser intensities were kept constant across tissues for the same fluorescence channel, but confocal exposure time and gain were adjusted according to each tissue to prevent signal saturation. Because fluorescence intensity scales linearly with exposure time and gain on our EMCCD, nuclear GFP/mScarlet-I levels between tissues in Figure 2 can be compared using these settings.

Images were all taken as single slices, with the exception of Figure 5A head images, which were taken as z-stacks spanning the head at a step size of 0.5 μm. The anatomical region imaged for each tissue was constant throughout the study, and are as follows: For intestinal cells, the anterior-most 2-4 nuclei were analyzed. For neurons, ventral nerve cord nuclei posterior to the vulva were analyzed. For muscle cells, nuclei at the dorsal side of the region between pharynx and vulva were analyzed. For hypodermal cells, hyp7 cell nuclei located at the lateral plane of the region between pharynx and vulva were analyzed.

### Image analysis for quantification of nuclear GFP/mScarlet-I and mScarlet-I levels

Confocal Image analysis was done using Fiji software^82^. First, background signals were subtracted from the GFP and mScarlet-I channel images. Individual cell nuclei were defined in the mScarlet-I channel image using the threshold function. These threshold-defined cell nuclei masks were applied independently to the GFP and mScarlet-I channel images to measure the mean GFP and mScarlet-I intensities in each cell nuclei. Mean GFP and mean mScarlet-I levels of each nuclei were divided to obtain the GFP/mScarlet-I value of that nuclei. For experiments where each n value is a single animal, a mean of the GFP/mScarlet-I values of all the nuclei imaged in that animal is reported. For experiments where each n value is a nuclei (specifically Figures 6 and S5), the GFP/mScarlet-I value of that nuclei is reported. For experiments where each n value is a single animal, the mean mScarlet-I value of all the nuclei in that animal was multiplied by the mean area of all the nuclei in that animal to obtain the nuclear mScarlet-I level. For experiments where each n value is a nuclei (specifically Figures 6 and S5), the mean mScarlet-I value of that nuclei is multiplied by the area of that nuclei to obtain the nuclear mScarlet-I level.

For Figures 5A and S4C, maximum z-projection projection images of the acquired z-stack images were used as input for this analysis. Threshold function defined a mask encompassing all the head nuclei instead of individual nuclei, which was used for measuring mean GFP and mean mScarlet-I levels of head nuclei of an animal.

### Aldicarb paralysis assay

18-25 L4 animals of each genotype were placed onto OP50-seeded NGM plates with aldicarb (100 mM stock solution, dissolved in 70% ethanol) added to a final concentration of 1mM. Every 10 minutes for a total of 2 hours, all animals on the plate were scored for paralysis, with paralysis defined as no movement of any kind in response to platinum wire worm pick prodding. Paralyzed animals were picked off the plate. Percentage of mobile animals at each time point assayed was plotted against duration. Three assay replicates were performed on separate days.

### Protein structure predictions

Protein interaction prediction in Figures S1A was generated by AlphaFold3 server^83^. Protein structure predictions in Figures 4A and S3A were generated by AlphaFold Protein Structure Database^58,59^. Protein structures were displayed and colored on The PyMOL Molecular Graphics System, Version 3.13.1 Schrödinger, LLC. pLDDT color scheme was developed by Konstantin Korotkov (https://github.com/sokrypton/ColabFold).

### Multiple sequence alignment

Protein sequences were imported from Uniprot^84^. Multiple sequence alignment was performed with MUSCLE v3.8.31 in Jalview v2.11.4.1^85–87^.

### Statistical analysis

Statistical analysis was done using Graphpad Prism for macOS, Boston, Massachusetts USA, www.graphpad.com.

## Author Contributions

Conceptualization, C.C., A.J.; Investigation, C.C.; Formal analysis, C.C.; Funding acquisition, C.C., A.J.; Writing - original draft, C.C.; Writing - review & editing, C.C, A.J..

## Acknowledgements

We thank members of the Jain lab, Nicolas Lehrbach, Peter Reddien, Fred Winston, and Iain Cheeseman for helpful discussions. We thank Stephen Nurrish and Josh Kaplan for the p*unc-25*::NLS::mTagBFP2 marker plasmid. We thank Christalyn Ausler for helping prepare NGM plates. We thank Steve Flavell, Yukiko Yamashita, and Scott Kennedy labs for kindly allowing us to use their lab equipment. This work is supported by grants from the NIH R35GM151111 (A.J.), Bumpus Foundation (A.J., C.C.), Pew Charitable Trusts (A.J.), and Chan Zuckerberg Initiative DAF2022-250422 (A.J.). C.C. is a William N. & Bernice E. Bumpus Fellow supported by the William N. & Bernice E. Bumpus Foundation.

## Resource Availability

Strains generated in this study are available upon request.

## Supplemental Information

Document S1. Figures S1–S5, Supplementary Note, Supp Note Fig 1, Supp Table 1.

