## Supplementary Information for "Polyamine homeostasis *in vivo* relies primarily on transport"

**Figures S1-S5**

**Supplementary Note**

**Supplementary Note Figure 1**

**References**

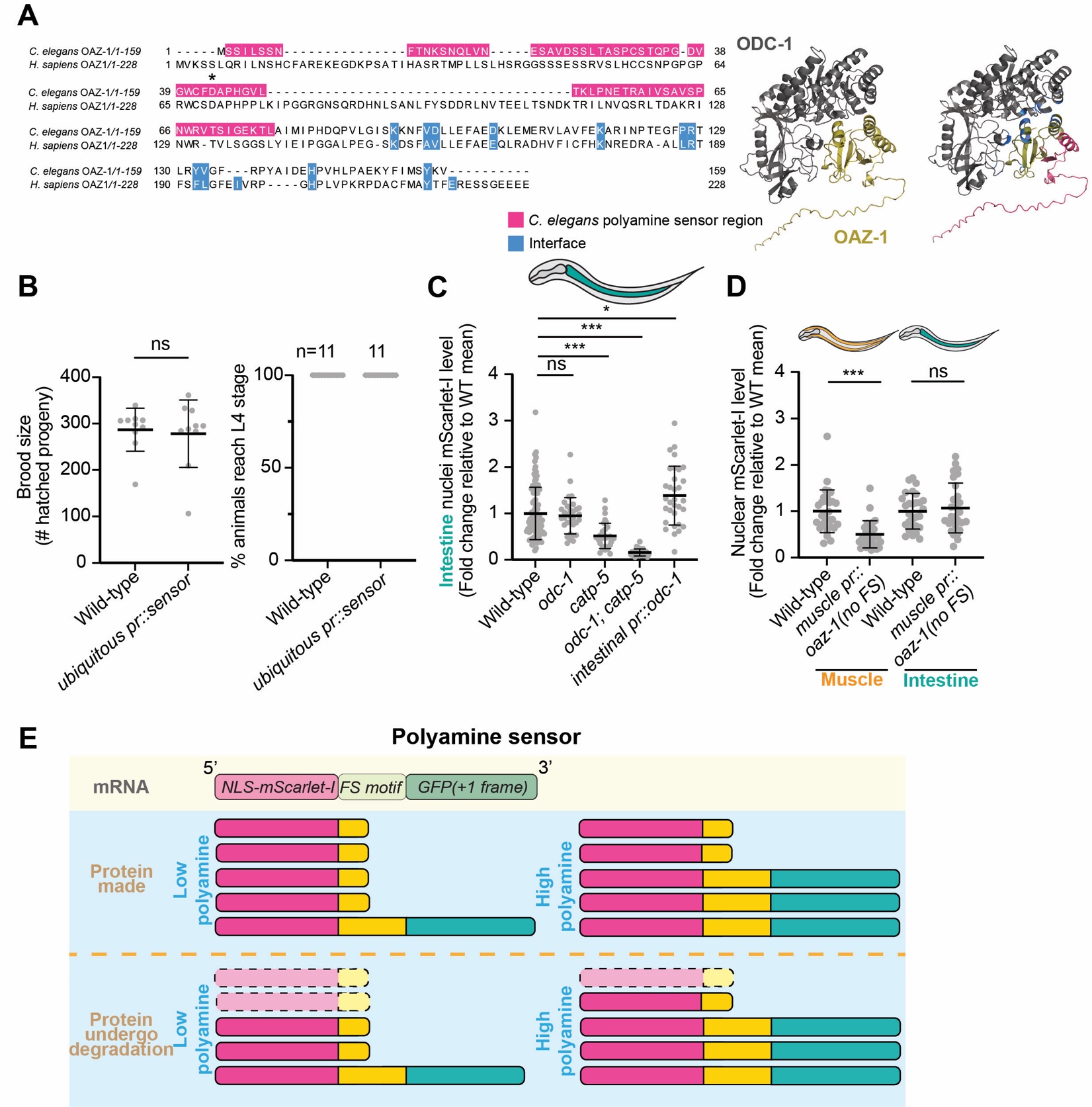

**Figure S1. (Related to Fig 1) Design and validation of a polyamine sensor in *C. elegans.*** (A) (Left) Protein sequence alignment of *C. elegans* OAZ-1 and *H. sapiens* OAZ1. Asterisk indicates frameshift site. Highlighted in pink: Residues in *C. elegans* polyamine sensor. Highlighted in blue: OAZ1 and corresponding OAZ-1 residues which form or strengthen the *H. sapiens* ODC and OAZ1 binding interface^1^. (Right) Alphafold3 predicted structure of *C. elegans* ODC-1 (dark grey) bound to OAZ-1 (gold) with (right structure) or without (left structure) residues highlighted. Highlighted pink residues are amino acids in OAZ-1 used in the polyamine sensor. Highlighted blue residues are amino acids in ODC-1 and OAZ-1 corresponding to human ODC1 and OAZ1 residues which form or strengthen the *H. sapiens* ODC-OAZ1 interface^1^. *H. sapiens*, *Homo sapiens*. (B) Animals expressing the ubiquitous promoter (p*eft-3*) driven polyamine sensor have normal brood size (n=10 per genotype across three biological replicates) (left) and development timing (right). Development timing was scored as percentage of progeny animals reaching L4 stage ~52 hours after egg stage. n indicates sample size (11 plates per genotype, with at least 20 progeny animals scored per plate). (C) Quantification of nuclear mScarlet-I in intestinal cells of animals with the indicated genotypes. Values are normalized to the mean value of wild-type animals. Corresponds to Figure 1B. (D) Quantification of nuclear mScarlet-I in muscle or intestinal cells expressing polyamine sensor. Values are normalized to the mean value of wild-type animals. Corresponds to Figure 1C. (B-D) Error bars denote mean ± SD. ns, not significant; *, p≤ 0.05; ***, p≤0.001 ((B,D) Mann-Whitney test (C) Kruskal-Wallis test with Dunn’s test for multiple comparisons). All experiments done with at least three biological replicates. (E) Model to explain that nuclear mScarlet-I level correlates with polyamine levels. Under the assumption that a constant fraction of NLS-mScarlet-I-only fragments is degraded, total nuclear mScarlet-I level (with contribution from both NLS-mScarlet-I-only and NLS-mScarlet-GFP proteins) will correlate with polyamine level. NLS-mScarlet-I-only proteins with dashed outline and faded coloring indicate degraded proteins. FS, frameshift.

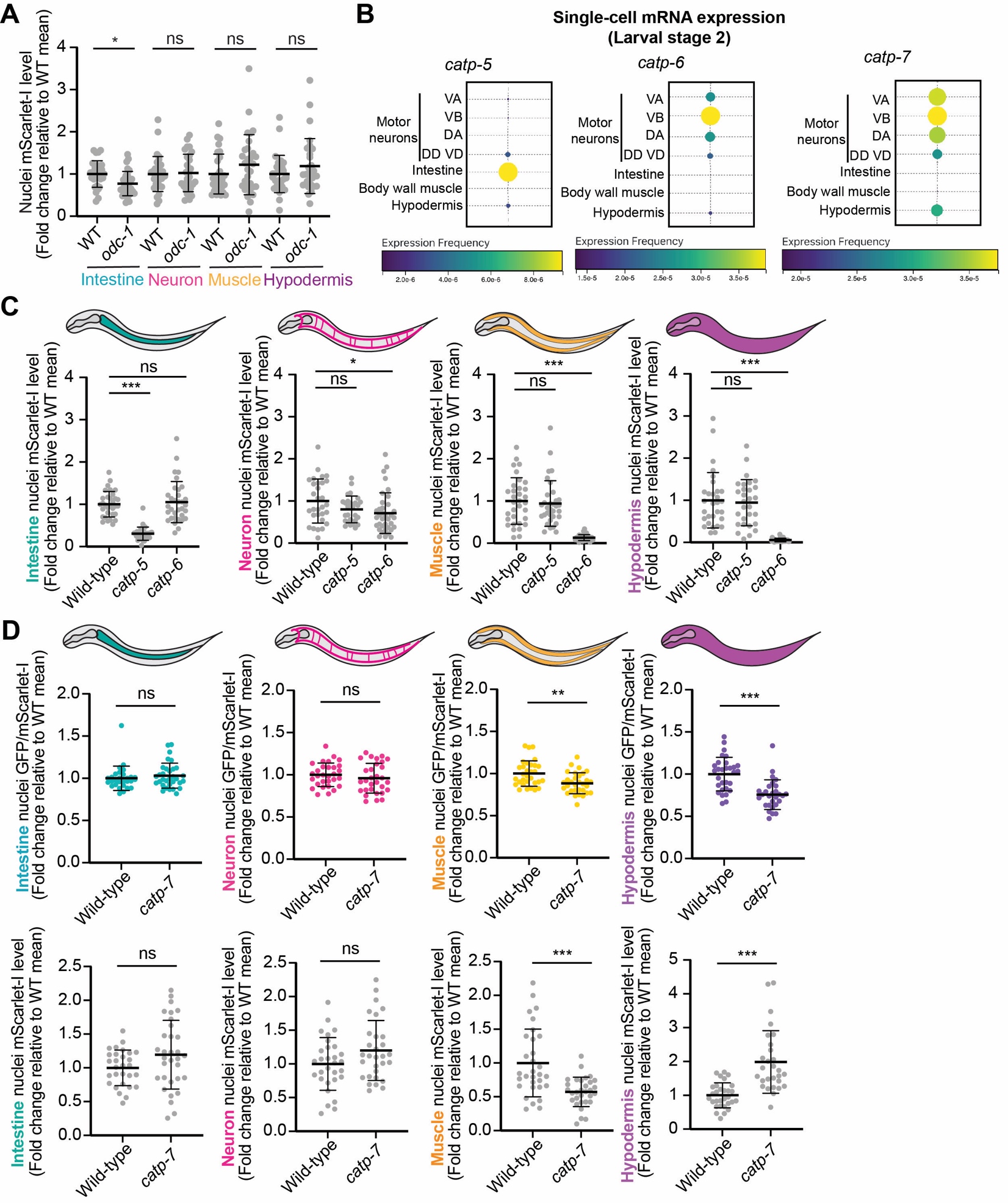

**Figure S2. (Related to Fig 3) Polyamine transporters are the primary modulators of polyamine levels in somatic tissues.** (A) Quantification of nuclear mScarlet-I in indicated tissues in wild-type and *odc-1* mutants. Values are normalized to the mean value of wild-type animals. Corresponds to Figure 3A. (B) Larval stage 2 single-cell mRNA expression levels of *catp-5*, *catp-6*, and *catp-7* across tissues. Data from Ben-David et al., visualized by WormBase visualization tools for *C. elegans* single-cell data^2,3^. Expression frequency indicated by both color and size of dot. (C) Quantification of nuclear mScarlet-I levels in indicated tissues in wild-type and polyamine transporter mutants. Values are normalized to the mean value of wild-type animals. Corresponds to Figures 3B and 3C. (D) Quantification of nuclear GFP/mScarlet-I (top) and nuclear mScarlet-I levels (bottom) in indicated tissues in wild-type and *catp-7* mutants. Values are normalized to the mean value of wild-type animals. Note that results for hypodermis are inconclusive due to nuclear GFP/mScarlet-I and mScarlet-I levels changing in opposite directions in WT and *catp-7* mutant animals. (A,C,D) n=27-36 animals per genotype across three biological replicates. ns, not significant; *, p≤ 0.05; **, p≤ 0.01; ***, p≤0.001 ((A,C) Kruskal-Wallis test with Dunn’s test for multiple comparison; (D) Mann-Whitney test). Error bars denote mean ± SD.

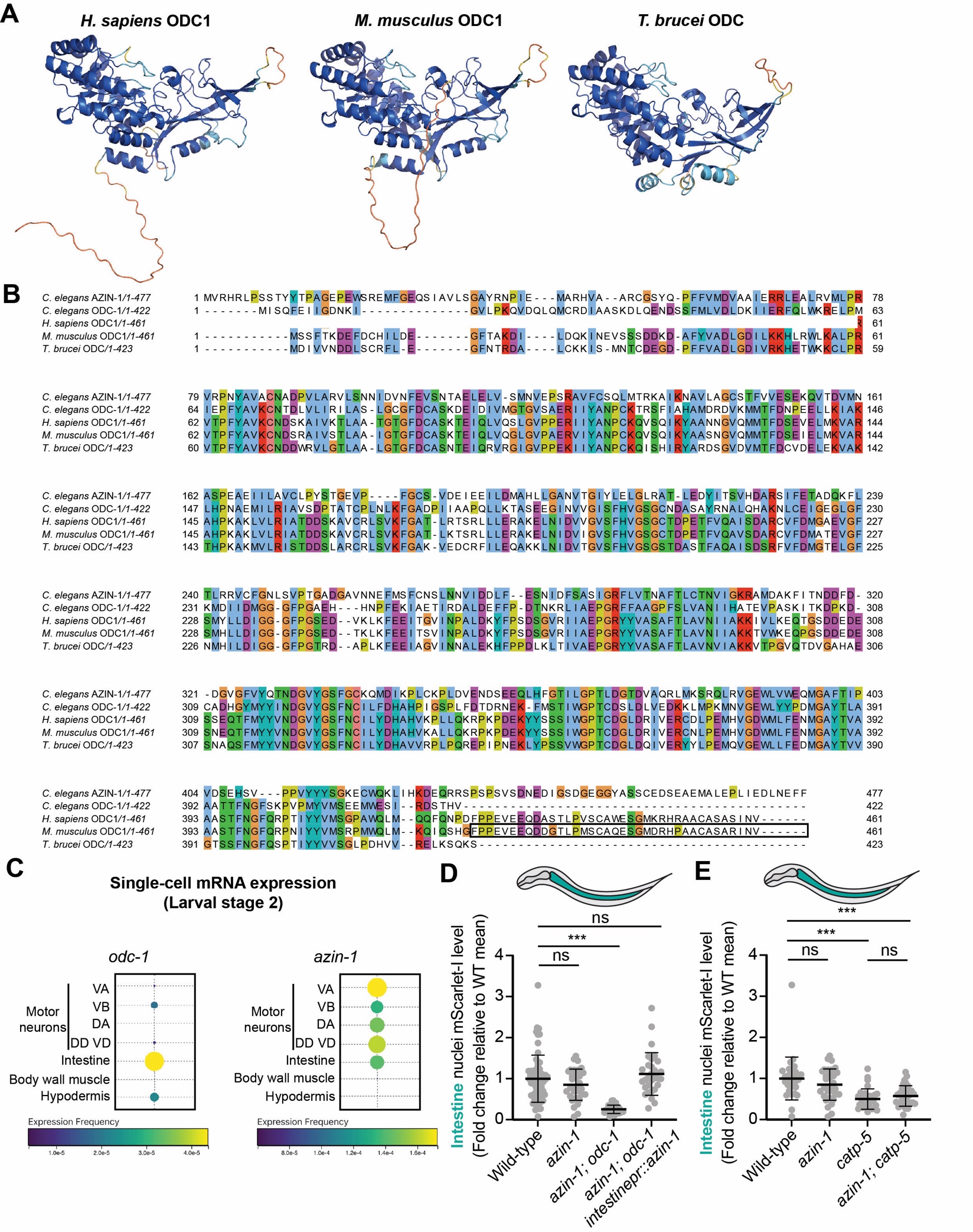

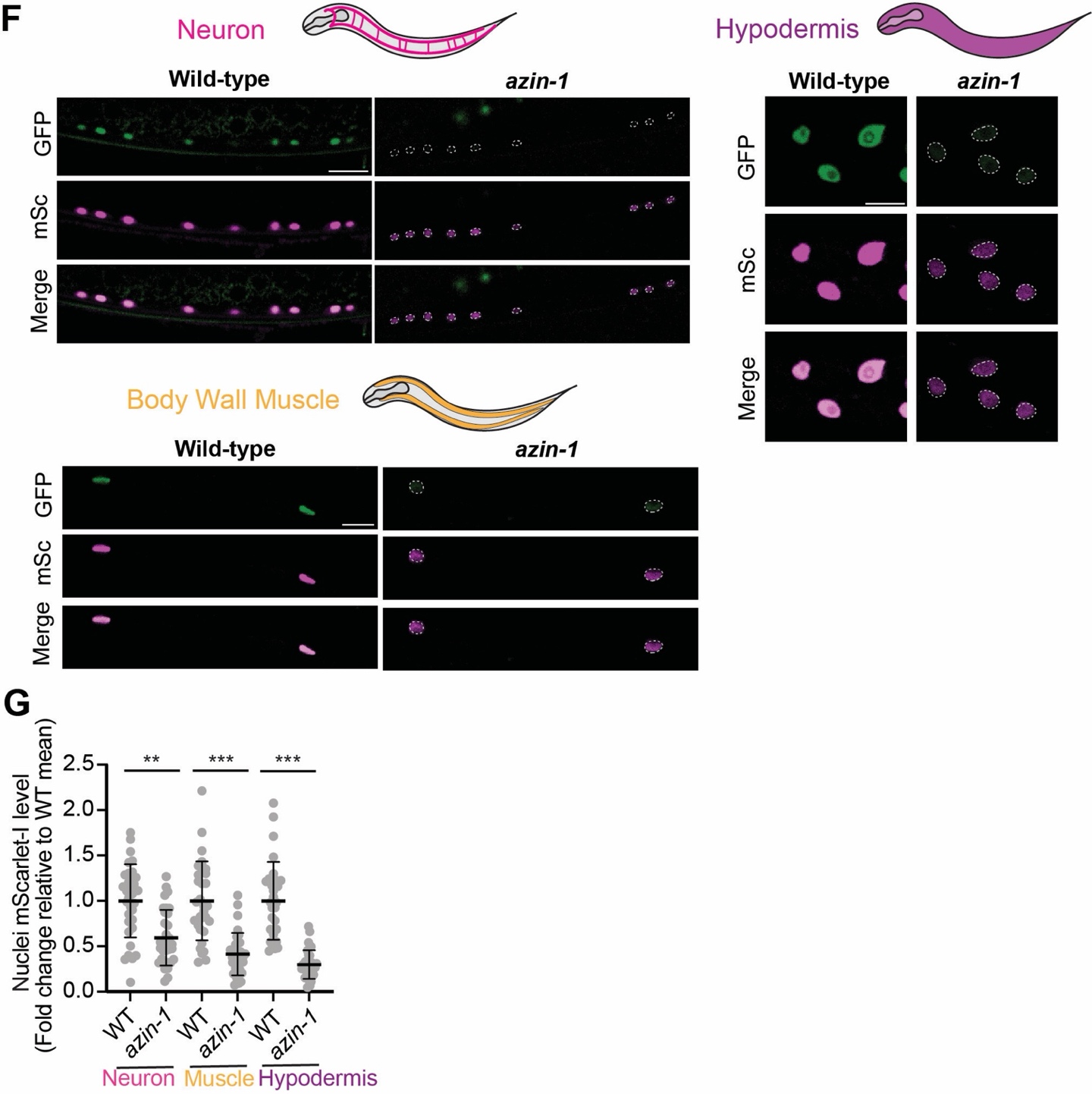

**Figure S3. (Related to Fig 4) AZIN-1 regulates polyamine transport.** (A) Alphafold2 structure predictions of ODC in various species^4,5^. Note the extended C-terminus tail region present in *H. sapiens* and *M. musculus* ODC1, but absent in *T. brucei* ODC. *M. musculus*, *Mus musculus*. *T. brucei*, *Trypanosoma brucei.* (B) Protein sequence alignment of *C. elegans* AZIN-1, *C. elegans* ODC-1, *H. sapiens* ODC1, *M musculus* ODC1, and *T. brucei* ODC1. The last 37 amino acids of mouse ODC1, sufficient for the short half-life of ODC1, are boxed. Amino acid colouring by Clustal X colour scheme. Figure generated by Jalview v2.11.4.1 using MUSCLE v3.8.31 algorithm^6–8^. (C) Larval stage 2 single-cell mRNA expression levels of *odc-1* and *azin-1* across tissues. Data from Ben-David et al., visualized by WormBase visualization tools for *C. elegans* single-cell data^2,3^. Expression frequency indicated by both color and size of dot. (D,E,G) Quantification of nuclear mScarlet-I levels in indicated tissues of indicated genotypes. Values are normalized to the mean value of wild-type animals. Corresponds to Figures 4B (D), 4C (E), and 4D (G). n=29-33 per genotype across at least three biological replicates. ns, not significant; **, p≤ 0.01; ***, p≤0.001 (Kruskal-Wallis test with Dunn’s test for multiple comparisons). Error bars denote mean ± SD. (F) Representative fluorescence images of neuron, muscle, and hypodermis nuclei expressing polyamine sensor in wild-type and *azin-1* mutant animals. mSc, mScarlet-I. Scale bar, 10 μm.

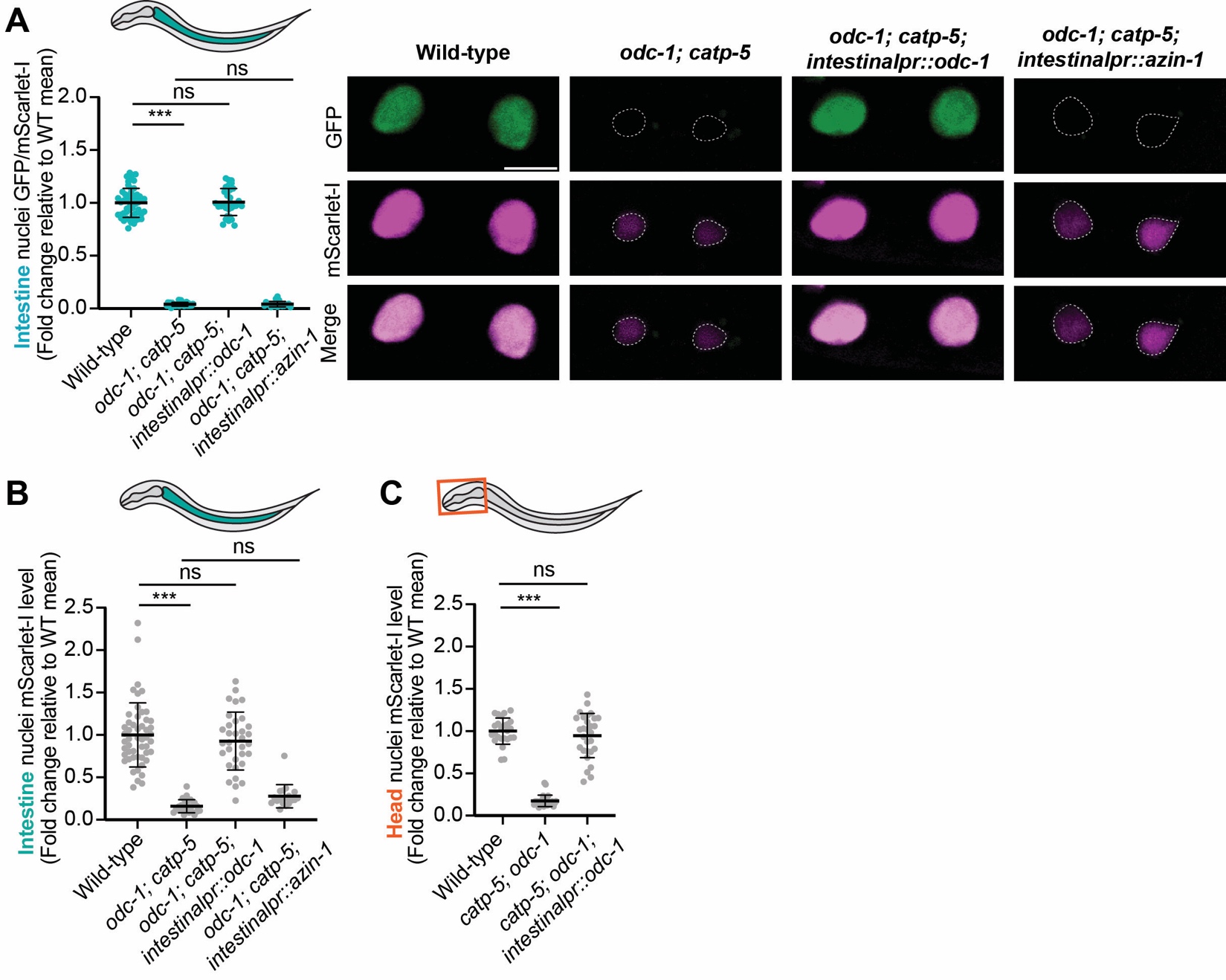

**Figure S4. (Related to Fig 5) The intestine is a central hub for organismal polyamine regulation.** (A) (Left) Quantification of intestinal polyamine levels in indicated genotypes. Values are normalized to the mean value of wild-type animals. (Right) Representative fluorescence images show corresponding sensor expression in intestinal nuclei. Scale bar, 10 μm. (B,C) Quantification of nuclear mScarlet-I levels in indicated tissues of indicated genotypes. Values are normalized to the mean value of wild-type animals. Corresponds to Figures S4A (B) and 5A (C). (A-C) n=20-52 per genotype across at least three biological replicates. ns, not significant; ***, p≤0.001 (Kruskal-Wallis test with Dunn’s test for multiple comparisons). Error bars denote mean ± SD.

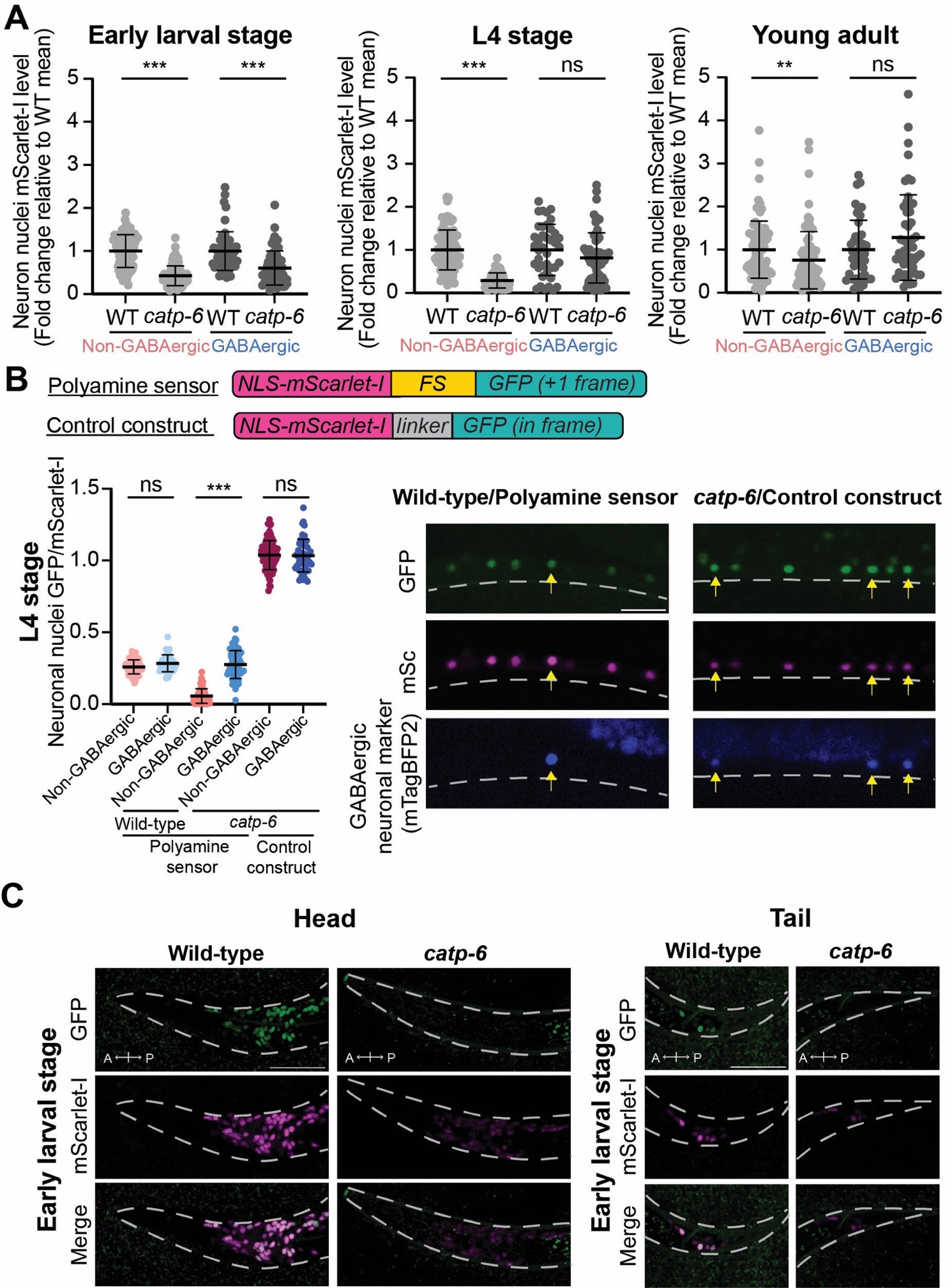

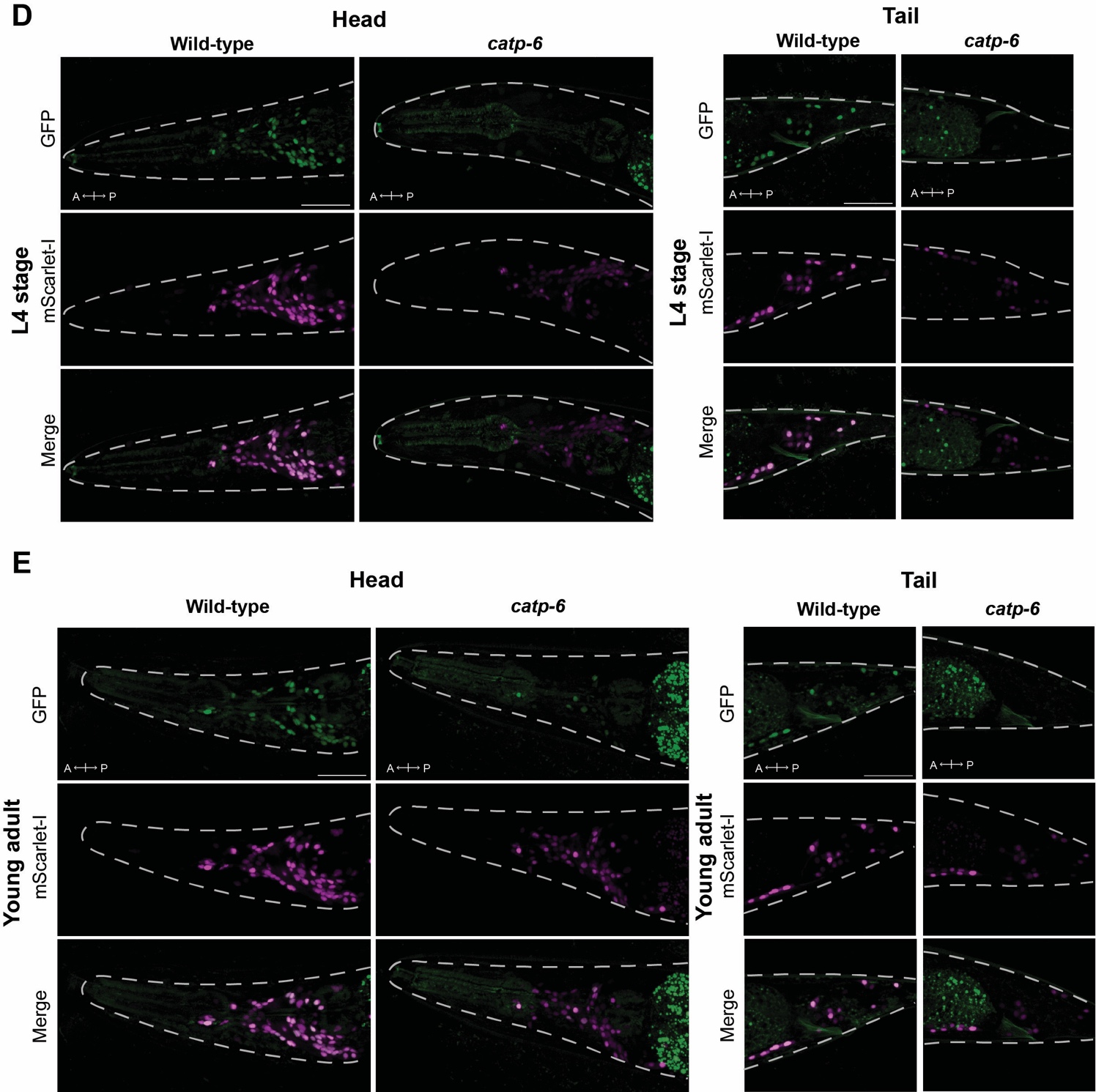

**Figure S5. (Related to Fig 6) Polyamine transporter dependency changes during nervous system development.** (A) Quantification of nuclear mScarlet-I levels in VNC non-GABAergic and GABAergic neurons in wild-type and *catp-6* mutants across developmental time points. Values are normalized to the mean value of the same neuron type and of the same developmental stage in wild-type animals. Corresponds to Figure 6B. (B) Control experiment showing that the L4 stage *catp-6*-dependent regulation of polyamine levels in GABAergic neurons measured by the polyamine sensor is not an artifact of different general translation and/or transcription efficiencies in non-GABAergic versus GABAergic neurons. (Left) Quantification of VNC neuronal nuclei GFP/mScarlet-I in L4 stage animals of indicated genotypes. (Right) Representative fluorescence images of VNC nuclei in indicated conditions. GABAergic neuronal nuclei are marked with mTagBFP2 and indicated with yellow arrows. Scale bar, 10 μm. FS, frameshift motif. (A,B) Each data point (n) is a neuronal nuclei GFP/mScarlet-I value. n= 40-94 from 30-35 animals for each condition (genotype/stage/neuron type combination) across at least three biological replicates. Error bars denote mean ± SD. ns, not significant; **, p≤ 0.01; ***, p≤0.001 ((A) Mann-Whitney test; (B) Kruskal-Wallis test with Dunn’s test for multiple comparisons). (C) Fluorescence images of head and tail region neuronal nuclei expressing polyamine sensor in WT and *catp-6* mutants during early larval stage, L4 stage, and young adult stage. Scale bar, 25 μm. A, anterior direction; P, posterior direction (of animal).

**Supplementary Note**

Polyamines are known to regulate translation efficiency, so reduced nuclear mScarlet-I levels under low polyamine conditions could also reflect decreased overall protein synthesis. However, the following experiment suggests that this is not the case: In L4 stage *catp-6* mutants, non-GABAergic neurons of the ventral nerve cord (VNC) have decreased polyamine levels, but not GABAergic neurons (Figure 6B). If a decrease in general translation is causing nuclear mScarlet-I levels to decrease under low polyamine conditions, we would expect nuclear mScarlet-I levels to be low in L4 stage *catp-6* mutants expressing either the polyamine sensor or a control GFP-linker-mScarlet-I construct. However, we only see this decrease in polyamine sensor-expressing animals and not control construct-expressing animals (Supplementary Note Figure 1).

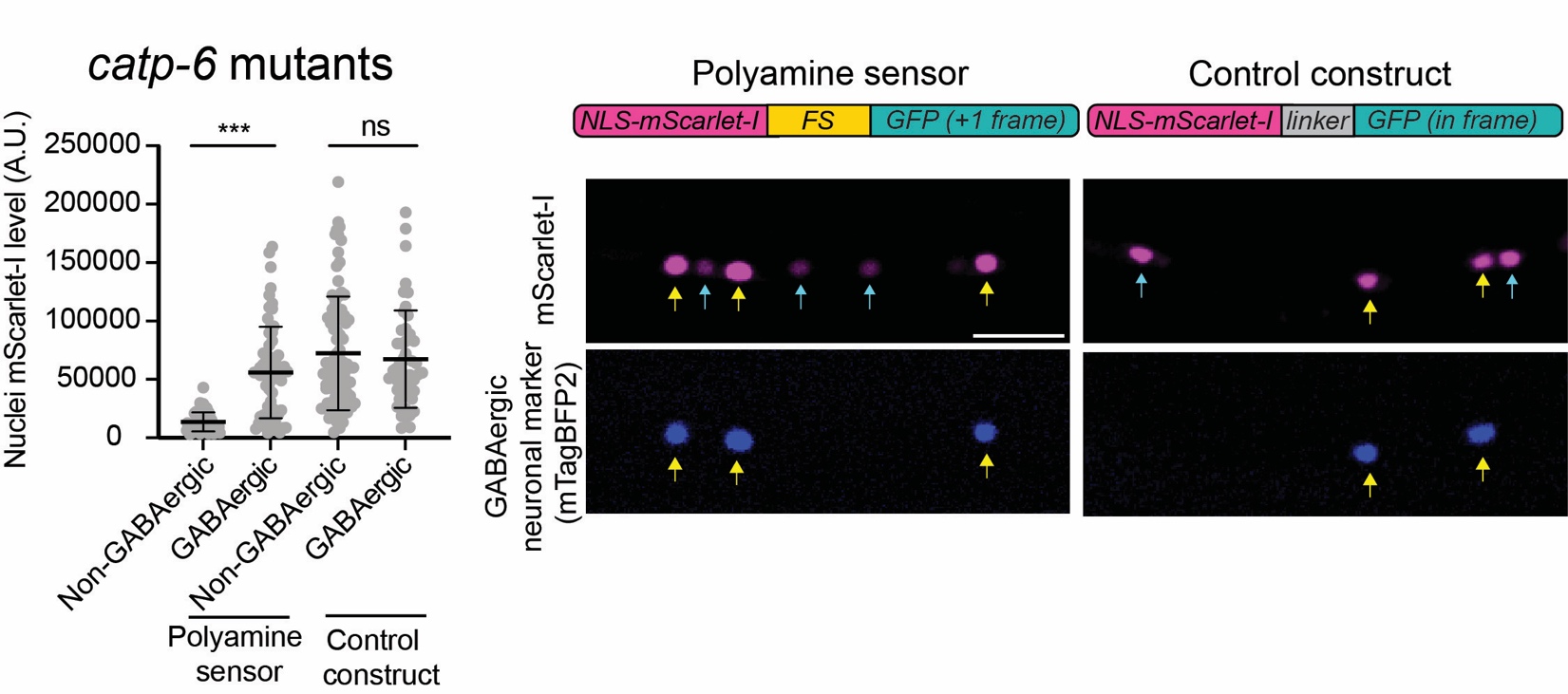

**Supplementary Note Figure 1.** (Left) Quantification of nuclear mScarlet-I levels in indicated neuron types in L4 stage *catp-6* mutants expressing either the polyamine sensor or the control construct. Each data point (n) is a neuronal nuclei GFP/mScarlet-I value. n= 50-91 from 30-31 animals for each condition (genotype/neuron type combination) across at least three biological replicates. Error bars denote mean ± SD. ns, not significant; ***, p≤0.001 (Mann Whitney test). (Right) Representative fluorescence images of nuclear mScarlet-I expression in VNC neurons of L4 stage *catp-6* mutants expressing either the polyamine sensor or the control construct. GABAergic neuron nuclei are labeled with mTagBFP2 (yellow arrows). Unlabeled neuronal nuclei are non-GABAergic neuron nuclei (blue arrows). Scale bar, 10 μm. A.U., arbitrary units.
